# Inclusive Biology Curriculum Interventions Can Reduce High School Students’ Bioessentialist Beliefs

**DOI:** 10.64898/2026.04.16.719004

**Authors:** Charlie K. Blake, Onyebuchi S. Ewa, Emily B. Eckles

## Abstract

Lesbian, gay, bisexual, transgender, queer, intersex, and asexual (LGBTQIA+) students continue to face violence, exclusion, and barriers at school, including in STEM education. A key underexamined factor in diversity, equity, and inclusion (DEI) efforts is the content of the life science curriculum, which is uniquely positioned to reinforce or refute bioessentialist, binary, and heteronormative biases. Outdated science curricula not only conflict with current scientific evidence but can also perpetuate beliefs that contribute to sexism and LGBTQIA+ marginalization. To address this, we designed four gender and sexual diversity (GSD)-inclusive biology activities, aligned with NGSS standards, and informed by inclusive curriculum frameworks. Using a mixed-methods approach, we studied 127 high school students who participated in two or more inclusive biology activities. Surveys conducted before and after implementation showed significant reductions in essential, binary beliefs about sex and gender, and increases in affirming attitudes toward sex and gender diversity. Interviews conducted after implementation further revealed differences between LGBTQIA+ and straight students’ conceptualizations of biological sex. Our findings demonstrate that even brief curriculum interventions can shift student attitudes, although we hope future studies will explore the impact of sustained interventions. Updating life science instruction is essential for educational equity and scientific accuracy.

## INTRODUCTION

While there has been increasing attention to diversity, equity, and inclusion (DEI) in K-12 schools in recent years, we also know that a pattern of social exclusion and violence persists for lesbian, gay, bisexual, transgender, queer, intersex, and asexual (LGBTQIA+) students. Recent school climate data indicates that over 31% of LGBTQIA+ students were pushed, shoved, or physically assaulted by peers at school due to their perceived sexual orientation, gender expression, or gender (Kosciw et al., 2022). There is also growing evidence that when it comes to the workforce, LGBTQIA+ people may represent the largest and most understudied group that is underrepresented in STEM fields (Freeman, 2020, 2021). These disparities in safe and equitable access to education and STEM careers affect a growing group of millions of students who claim an identity somewhere under the LGBTQIA+ umbrella, over 20% of the generation recently or currently in high school (Gallup, 2024; IPSOS, 2021).

In an attempt to address the disparities facing the LGBTQIA+ student population, many schools focus their interventions on important strategies like staff trainings, student counseling, and institutional policy changes—with less attention paid to equally important critical pedagogical examination of curriculum within disciplinary courses (Corsino & Fuller, 2021; Freire, 2000; IWG on Inclusion in STEM, 2021; Wright & Delgado, 2023). Only about 2% of LGBTQIA+ students in recent years reported seeing gender and sexual diversity (GSD) inclusive curriculum in their science courses (Kosciw et al., 2022). Yet research has shown that representation in the curriculum is associated with reduction in anti-LGBTQIA+ violence and increased likelihood of choosing a STEM major for LGBTQIA+ high school students (IPSOS, 2021; Kosciw et al., 2013, 2014), and increased performance for undergraduate students (Kelly et al., 2022). Research has also shown that students at the undergraduate level are not only capable of noticing normative ideologies in outdated textbook content, but also suggesting corrections for improved, scientifically precise language (Dunk et al., 2024). Thus, addressing LGBTQIA+ representation and equity in science course content should be of great concern if we hope for equitable access to education for all, and a diverse and inclusive scientific community going forward.

Rather than being based on the most up-to-date scientific evidence that indicates the biology of sex is complex and nuanced, many life science topics are currently taught using outdated content that has been influenced by sociocultural biases (Bazzul & Sykes, 2011; Snyder & Broadway, 2004; Zemenick et al., 2022). We will describe three common types of biases here: sex and gender bioessentialism, sex and gender binarism, and heterosexism or heteronormativity. One type of bias is biological essentialism or bioessentialism, which overstates the importance of genetics and biological processes in determining phenotypic and behavioral characteristics, and erroneously links human social norms to supposed underlying deterministic biological causes (Dar-Nimrod & Heine, 2011; de Melo-Martín, 2003; Haslam et al., 2006). The influence of essentialist thought across multiple scientific disciplines stems from deep foundational roots, such as the highly influential work of Konrad Lorenz, who popularized the idea of innate instincts, and was also a eugenicist and member of the Nazi party (Blumberg, 2005; Lehrman, 1953). Bioessentialism and deterministic thinking often rely on taking subjectively perceived sociocultural generalizations and constructing a pseudoscientific explanation for why they are inevitable, natural, right, and/or unchangeable. Essentialism may focus on the natural kind or essence of things (Neufeld, 2022), and bioessentialism may incorporate a specific link to biology or genetics as the source of determining these inevitable, essential traits (Harden, 2023). Essentialism can encompass many related patterns of thinking such as genetic determinism, and the natural immutability of race and ethnicity, but the current study will focus mostly on sex and gender essentialism that ascribes deep and deterministic differences on the basis of sex and/or gender (Greene, 2020).

Another type of bias, binarism, contrives dualistic and strictly dichotomous categories as a biological rule. Sex is often presented falsely as an easily defined and binarily categorical attribute of individuals, despite intersex individuals existing across many species and in many variations (Fausto-Sterling, 1993; Fernandez-Garrido, 2022; Roughgarden, 2013), and the difficulties of pinning down universally applicable definitions of sex to begin with (Gorelick et al., 2017; McLaughlin et al., 2023; Warkentin et al. 2026). There are also numerous examples of species with intrasexual polymorphisms characterized by wide variation or multiple types within sexes of physical and behavioral traits, often paired with multiple distinct mating strategies that are an important part of the ecology of the species (Bagemihl, 1999; McLaughlin et al., 2023; Roughgarden et al., 2015). Binarism can slow and erode the progress of scientific research, and has led to biased and myopic views of the “sex chromosomes” and their roles unrelated to sexual development (Richardson, 2012; Richardson, 2013). Dualistic thinking itself can also promote hierarchy; as soon as there are two named groups, distinguishing attributes are ascribed to these two groups, and these categories can easily become hierarchical and self-perpetuating (Saguy et al., 2021). For example, fixed universal differences in sexual strategies between promiscuous male and coy female individuals have been largely debunked by the scientific community, but science education has taken a long time to catch up with these advances, and some science education materials still treat Darwin’s and Batemans’s ideas about sexual selection with underexamined loyalty (Fromonteil et al., 2023; Maier & Belk, 2010; Tang-Martinez, 2017).

Relatedly, science curricula may also contain heterosexist or heteronormative bias, which proffers heterosexual reproductive activity as the sole biologically and evolutionarily relevant phenomenon to the exclusion of many other vitally important sexual and social behaviors (Bagemihl, 1999; Bailey & Zuk, 2009; Bazzul & Sykes, 2011; Poiani, 2010; Sommer & Vasey, 2006). In actuality, many other traits and behaviors in a variety of contexts contribute to the overall lifetime fitness of an individual. Further, heterosexist bias has led to homosexual behavior being often ignored by science, although several evolutionary origins and possible contributions to fitness of homosexual behavior have been proposed (Monk et al., 2019).

Importantly, some have argued that no evolutionary “solution” is needed to justify the existence of such a widespread, well-documented phenomenon as homosexual behavior, and that demanding a suitable explanation veers into the dangerous territory of pan-adaptationism in which every observable trait is presumed to be an optimal target outcome reached through a direct process of natural selection (Bagemihl, 2000; Gould et al., 1979; Roughgarden et al., 2006; Stern et al., 2022). Heterosexist bias and reliance on outmoded ideas about sexual selection has also led to many aspects of female behavior, and females in general, being ignored or understudied (Ah-King, 2022). For example, heterosexist bias has contributed to the harmful and problematic assumption that reproduction and fecundity are the only important indicator of health and success, especially for females (Packer & Lambert, 2022; Roughgarden, 2013).

In addition to being ignorant of current scientific advances, outdated science content can be co-opted to shore up social injustices, including racism, homophobia, and misogyny (Cheung et al., 2021; Heyman & Giles, 2006; Longino, 1996; Prentice & Miller, 2006; Yaylacı et al., 2021). Further, in research on adults’ attitudes about gender identity and transgender issues in society, many people indicate that their attitudes about these issues are influenced by their understanding of science (K. Parker et al., 2022). Research has also shown that gender essentialist beliefs predict prejudiced attitudes against gender nonconformity even in young children (Fine et al., 2024), and that endorsement of heteronormative attitudes and beliefs is associated with engaging in explicitly intolerant behaviors like bullying people perceived to be dressing or acting like the opposite sex (Duncan et al., 2019). More recently, pseudoscientific falsehoods have been incorporated into repressive legislative policies targeting LGBTQIA+ people, especially transgender, nonbinary, and intersex people, in attempt to justify these policies with the supposed authority of science (Executive Order 14168, 2025). Considering these direct invocations of scientific authority to do harm, science educators and researchers therefore have a responsibility to engage with the relationship between science education and its sociopolitical context (Sedlacek et al. 2026).

### Theoretical Frameworks

We began with a critical pedagogical perspective and the idea that decisions about curriculum content and pedagogical methods are never neutral in their origins or impacts (Freire, 2000). We approach any curriculum as a reflection and reproduction of cultural values and social norms that may be implicitly or explicitly conveyed through curricular decisions (Szalacha, 2004; Wright & Delgado, 2023). A critical approach to educational interventions contrasts with other types of DEI interventions that focus on school safety or on equity through promoting fair treatment (though these approaches can also have benefits). Rather, a critical approach focuses on interrogating the cultural values and social norms that are implicitly or explicitly conveyed through pedagogical and curricular decisions (Szalacha, 2004; Wright & Delgado, 2023). By beginning with what the curriculum teaches about the underlying concepts that circumscribe the social norms that may lead to disparities in the first place, critical curriculum interventions have the power not only to change the school climate and student experiences, but also potentially shift the underlying assumptions on which social hierarchies are built and carried into the wider world. The life science curriculum is uniquely positioned to either uphold or refute beliefs about fundamental concepts like biological sex, what it means to be female, or what is “natural” human anatomy and behavior. Informed by critical pedagogical theory, we began this study with the idea that science educators can and should craft science curricula to combat sociocultural bias that is rooted in outdated science or pseudoscientific ideas.

Our focus on high school students is also important in this regard, because we see mandatory high school biology classes as key spaces where cultural beliefs about sex and gender are enacted, refined, or refuted, and these beliefs are subsequently carried back out of the classroom into the next generation of society. We also drew from some tenets of applied action research, working with practicing high school teachers to inform our curriculum development and striving to make our research relevant to pressing issues currently faced by practitioners in real situations (Avison et al., 1999; Coghlan & Brydon-Miller, 2014). Thus, we considered the creation of practical, classroom-tested curriculum products, useable by high school teachers beyond those in this study as an objective equal in importance to our research goals.

Scholarship around inclusive science education has proliferated in recent years and there are a variety of frameworks and recommendations we could draw from in approaching curriculum development within this study (Casper et al., 2025; Hales, 2020; Kean, 2021; Wright & Delgado, 2023; Zemenick et al., 2022). In developing this study we chose to focus on Long et al.’s (2019; 2021) Inclusive Curriculum Framework because it was developed by practicing high school teachers (some of our past and current collaborators) within the practical contexts of high school science classrooms. Long et al. (2021) define five key characteristics of inclusive curriculum. *Authenticity* acknowledges diversity while maintaining scientific precision.

*Continuity* starts with a diversity lens throughout instead of an isolated token lesson or staring with oversimplification that must later be contradicted. *Affirmation* normalizes diverse phenomena and frames these with curiosity rather than pathologization. *Anti-Oppression* actively encourages students to recognize patterns of injustice in society. *Student Agency* provides students opportunities for choice and incorporates student feedback into lessons as learning unfolds. Similarly, Rende & Johnson’s (2024) framework draws from experiences of transgender science educators, and calls attention to interrogating power, resisting essentialism, and embracing experiential knowledge as key elements of inclusive science pedagogy.

We also referred to Cooper et al’s 2020 recommendations for inclusive teaching, which were more focused on university learning environments, but are still relevant to the high school context. The most applicable aspects of these recommendations for this project include being thoughtful about the language used to describe the LGBTQIA+ community, with awareness that preferred language varies between individuals within the community and can change quickly. Cooper et al (2020) also offer several recommendations under the category of inclusive biology classrooms, such as discussing the full range of gender and sexuality within the biology curriculum and actively working to dispel myths about the biology of sex, reproduction, hormones, and genetics.

Existing inclusive education frameworks give many practical suggestions for how educators might remove stigmatizing content from curricula and incorporate inclusive and expansive perspectives. Long et al’s (2021) and Cooper et al’s (2020) inclusive curriculum frameworks were informed by scholarship on DEI as well as the personal experiences of the authors, many of whom belong to the LGBTQIA+ community themselves in addition to their credentials as scientists and science educators. As such, we place trust in these recommendations, and our goal is not to test the merits of calls for inclusion that come from experts in impacted communities. Rather, our hope in this study was to examine how the use of inclusive curriculum updates affect the attitudes and beliefs of students who experience them.

The current study builds from earlier research that has established the link between sociocultural biases and educational science content. Previous research has examined how exposure to content promoting or refuting biological essentialism can impact attitudes and beliefs around sex and gender, however many of these studies have been conducted with undergraduate participants (Brescoll et al., 2013; Keller, 2005). Similarly, Beatty et al.’s (2021) study exposed undergraduate students to an ideologically aware curriculum intervention and explored the impacts of this curriculum intervention. Brian Donovan and colleagues have explored how curricular content can affect K-12 student beliefs about gender and race, especially within the context of genetics and genomics (Donovan, 2016, 2017; Donovan, Semmens, et al., 2019; Donovan, Stuhlsatz, et al., 2019; Donovan et al., 2020). While Donovan et al. (2024; 2019) have examined the impact of bioessentialist messages in K-12 science education contexts, they focused their efforts on understanding the role of biased science curriculum content in perpetuating sexism that impacts girls, rather than exploring equally important impacts on bias that affects the LGBTQIA+ community. Further, many of the studies above use content interventions designed for a controlled experimental implementation rather than content designed to become curriculum resources useable beyond the scope of the experiment by practicing educators in a classroom context. Thus, there remains a strong need for empirical studies that seek to understand what impacts applied inclusive curriculum reforms may have on students, and through what mechanisms, especially curriculum approaches developed with and for practicing teachers at the high school level, and specifically designed to address inclusivity of sex and gender diversity.

### The Current Study

Given that the large and growing LGTBQIA+ student population faces pronounced educational barriers, disparities in STEM pathways, and social exclusion that can be informed by biases promoted by obsolete and misleading STEM curricula, and that curricular choices also impact sociocultural beliefs that all students take out into society, we propose carefully designed changes to the life science curriculum to ameliorate these issues. The present study describes the implementation and testing of a GSD-inclusive biology curriculum intervention through a quasi-experimental, longitudinal, mixed-methods approach working with high school teacher partners and high school student participants. Weaving together qualitative and quantitative findings simultaneously, we sought to understand how inclusive updates to the biology curriculum impact students through the following research questions:

RQ1. Does exposure to the curriculum intervention change the prevalence of heteronormative behavior beliefs, essential and binary gender beliefs, transphobia, or (in contrast) gender and sex diversity affirming attitudes and beliefs held by high school students, and what factors are important in shaping these student attitudes and beliefs?

RQ2. Do LGBTQIA+ high school students and their straight peers differ in their heteronormative behavior beliefs, essential and binary gender beliefs, transphobia, or gender and sex diversity affirming attitudes and beliefs, either before or after the curriculum intervention, and what important factors and experiences in science education relate to any differences in these groups attitudes and beliefs?

## METHODS

To answer the first part of RQ1 we compared quantitative survey scores before and after the intervention, and to address the second part we used qualitative interviews to probe what factors may be behind the quantitative trends we observed. Similarly, to answer RQ2 we examined differences in survey scores among demographic groups both before and after the intervention, and delved into qualitative data to understand what factors and experiences informed any differences in these attitudes and beliefs between groups. We note the we have decided to use the term LGBTQIA+ as the default term for this community in our study materials and in this manuscript. None of the options for terminology feel ideal, and this acronym seemed like the best compromise of considerations of expansiveness and practicality. Some of our author team use the term queer to describe ourselves, and when work that we reference, or students that we interviewed used other terms, we matched our language to theirs.

### Author Positionality

In an effort to cultivate trust and transparency with each other and with our readers, the research team reflected on how our identities and experience impact our work as researchers (Secules et al., 2021). The lead author and principal investigator of the project (CB) is a white, queer, nonbinary trans person who also identifies with the term third gender. Their experiences as a former student include taking AP science courses in high school and studying biology and ecology at the undergraduate and doctoral level. In the past, they have also conducted research as a behavioral ecologist, and have taught biology, ecology, environmental science, and environmental health at the university level and environmental education at the K-12 level. These experiences allowed the author to relate to both science students and science teachers in the study, as well as to scientists who create and shape current scientific knowledge. In addition, their cultural experiences within transgender and queer communities give them a nuanced understanding of many different ways that humans may understand and define sex and gender diversity. Their understanding of sex and gender is also informed by previously spending time in radical feminist spaces as a young adult. These experiences, in addition to knowledge gained from existing scholarship, shaped the way the author designed the study and interpretated data. For example, it was particularly important to this author to strive for demographics questions that would work well for queer and trans students since they have had so many personal experiences of alienation and erasure with demographic survey questions themselves. The author’s positionality also influences their understanding of terms like sex and gender, and their heightened attunement to interviewee’s varied interpretations of these terms in responding to interview prompts. It is also important to note that CB conducted the interviews online with their own camera turned on, but did not meet any of the study participants in person, which means students had access to some visual information about the sociocultural identities of the interviewer, but leaves it unclear what conclusions participants may have drawn about their interviewer that could have influenced their responses.

EE is a white, queer woman whose undergraduate and graduate studies explore the complexity of sexuality and gender in modern German history. History, like many STEM disciplines, is plagued by underrepresentation of women and LGBTQIA+ scholars and scholarship, and this has given the author a deep understanding of how bias may play out within an academic discipline. Her experiences as an early career historian, combined with her former experience in biology courses at the high school and undergraduate level in the state of Illinois, gave her a valuable perspective on the project from outside the STEM discipline.

OE is a Black woman pursuing graduate studies in Diversity and Equity in Education. She entered this research with a strong awareness of how one’s identity can shape educational experiences and a deep understanding of ways structural inequities are embedded into curriculum and pedagogy. OE held the position of both a student and investigator during the time of this research, which provided a dual perspective of studying and navigating inequities within the social and institutional structures in education. This perspective provided this author with a heightened sensitivity to how a misrepresentation of minority students, including LGBTQIA+ students, is affected when course content fails to reflect current scientific understanding. While OE is not personally identified as LGBTQIA+, through this project, she supports the challenges to the hidden biases in biology curriculum in order to contribute to a more inclusive representation of sex, gender, and sexuality in STEM education.

The research team also notes that the funding for this project was targeted for termination due to its content, beginning with the listing of this study in an October 2024 investigative report released by the U.S. Senate Committee on Science, Commerce, & Transportation. Ultimately funding for the project was removed the following year and authors OE and EE lost their employment due to the removal of funding. The threat of, and eventual loss of funding affected the psychological well-being of the researchers over the course of the study, raised the personal and professional stakes of the study for the researchers, and shortened the originally planned study timeline. Teacher partners were made aware of the funding termination but still received their promised stipends, and student participants were not aware the study was targeted for funding termination.

### Curriculum

To create our curriculum intervention, we developed four GSD-inclusive biology activities designed to address common issues in the traditional life sciences curriculum. We began by reviewing the content of several biology textbooks commonly used in US high schools to identify topics particularly in need of updating, similar to methods used by previous researchers who have found science textbooks contain significant biased content (Bazzul & Sykes, 2011; Donovan et al., 2024). As we designed the updated materials, we ensured that all activities were Next Generation Science Standard (NGSS) aligned and focused on learning objectives that are priorities for high school life science educators. The materials were designed with Sam Long et al’s (2021) Inclusive Curriculum Framework as a guide and our approach to curriculum development also aligned with guidelines from Cooper et al., (2020) that recommend an inclusive science curriculum discuss the full range of gender and sexuality in biology class and avoid assuming social relationships and social identities in favor of more precise terminology focused on biological components and processes.

For example, the development of one of our biology acitivites began with examing the Reproduction lesson in a widely-used biology textbook, which features a diagram of the life cycle of a jellyfish (Miller & Levine, 2019, p. 882). The content of this section is meant to highlight an organism with more than one body type and a complex life cycle that involves both sexual and asexual reproduction in several different reproductive body forms. Our modified lesson focuses on the same themes but replaces the jellyfish example with a symbion (*Symbion pandora*, Figure 1). Like jellyfish, symbions are marine organisms with complex life cycles involving both sexual and asexual phases (Funch & Møbjerg Kristensen, 1995; Neves et al., 2010). However, the life cycle of symbions contains more than two reproductive forms that cannot be described solely with the terms male and female. The modified content describes the multiple body types and sexual roles present in this organism, conveying the same learning objectives about evolution of reproductive strategies as the original reading, but avoiding the assumption that there can only be two dualistic roles in sexually reproducing organisms.

**FIGURE 1.**
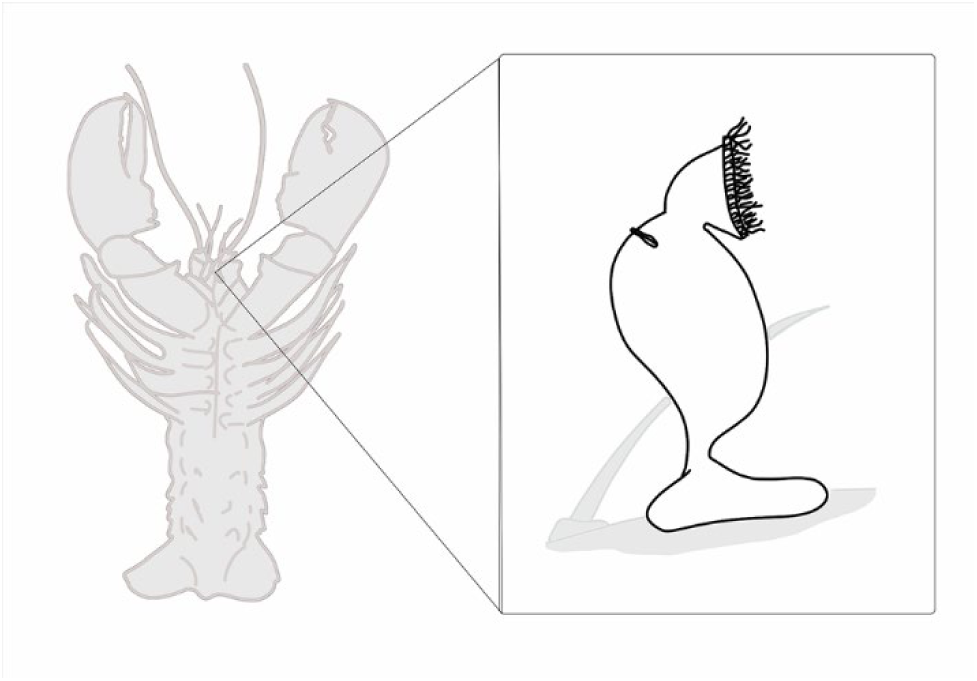
The pandora form of the *Symbion*, a small organism that lives on the mouthparts of lobster. This illustration is a sample of content from the inclusive biology activities.

Similarly, our lessons on animal bodies and behavior used examples of species with frequent occurrence of intersex individuals or same-sex parental pairs to explore factors that affect the evolution of mating systems and reproductive behaviors.

We took an analogous approach to develop each of the activities, using the most current science to update topics commonly taught in high school science classrooms. While our research questions are focused on how the curriculum intervention affected attitudes and beliefs around sex and gender diversity, we incorporated related inclusive changes into our lessons when relevant, such as replacing the term “pedigree” with “genetic inheritance chart” due the ableist and eugenicist history of the older term (Hales, 2020; Shotwell, 2021). This curriculum development process resulted in four activities, 20-60 minutes each, covering reproductive life cycles, genetic inheritance, animal bodies and behavior, and sex determination and development (visit our website for the full curriculum https://stemcenter.siue.edu/). The curriculum went through multiple iterative rounds of review, incorporating feedback from educators with expertise in inclusive teaching, and high school teachers who implemented the activities in their classrooms with diverse student populations (Table 1).

**Table 1.** Timeline and process of curriculum development and data collection

| Curriculum development |  |  |  |  |
| --- | --- | --- | --- | --- |
| Summer 2023 | Fall 2023 | Summer 2024 | Summer 2025 | Fall 2025 |
| Explored inclusive teaching frameworks and existing biology textbooks to identify key curriculum topics. Developed 4 draft lessons. | Lessons reviewed by inclusive teacher experts and feedback incorporated. | Reviewed by consulting education scholar.<br>Edited teacher materials and minor edits to lesson plans based on teacher and student feedback. | Third implementation by the authors in informal learning environment for iterative curriculum improvement only. | Incorporated student and teacher feedback.<br>Added supplemental PowerPoint files to curriculum.<br>Curriculum shared publicly. |
| Implementation and data collection |  |  |  |  |
|  | Spring 2024 | Spring 2025 | Fall 2025 |  |
|  | First implementation by classroom teachers | Second implementation by classroom teachers and online<br>Participant interviews | Follow up survey |  |

### Data collection

In the spring semesters of 2024 and of 2025, we surveyed participants within two weeks before and within two weeks after their participation in the curriculum intervention, and all participants were also offered the opportunity to participate in an interview two to six weeks after participating in the activities. Online participants were also offered the chance to complete follow-up surveys four to six months after their initial participation. Each implementation of the activities differed based on student and teacher needs, but each student completed at least two of the inclusive activities. Students who were 18 consented to participation for themselves, and students who were under 18 provided their informed assent along with parental informed consent. The protocol for this study was approved by the Institutional Review Board of Southern Illinois University Edwardsville (Protocol #2171).

### Study instruments

We used previously validated scales to measure student attitudes and beliefs. We used both of the subscales from the Heteronormative Attitudes and Beliefs survey (HABS), each containing 8 items, some of which we reverse scored as indicated by the creators (Habarth, 2015). In our study we term these two subscales the Heteronormative beliefs: normative behavior scale, which focuses on attitudes about how people should behave based on their role, and the Heteronormative beliefs: essential, binary sex and gender scale, which focuses on beliefs about the degree to which gender is fixed, distinct, and binary. Our Transphobia scale comes from Nagoshi et al. (2008), which assesses attitudes of fear and prejudice towards individuals who do not conform to socially expected gender roles. The HABS and Transphobia scales were both developed with undergraduate students with mean age 19-20 (Habarth, 2015; Nagoshi et al., 2008), supporting validity for use with our similar study population of high school students, the majority of whom were 18. From the Gender/Sex Diversity Beliefs Scale (GSDBS) we used the affirmation subscale (Schudson & van Anders, 2022), which captures attitudes that affirm the existence of diverse sexual and gender identities as well as beliefs about the fundamental nature of sex and gender. Of the 14 original questions in the affirmation subscale we removed one question, “Transgender people were born the way they are,” because this perspective remains controversial within transgender communities (Faye, 2018). The GSDBS was originally developed and validated with a wide age range of participants, with the original study population including 18-year-old participants and participants who had not completed high school (Schudson & van Anders, 2022). We retained the original format of response options for each scale, so the transphobia scale offered six Likert response options, and the other scales offered seven Likert response options. We also note that our survey included a scale that was later discarded due to unreliability and not included in the findings, the 11-item science subscale of the STEM Career Interest Survey (STEM-CIS), which was intended to measure students’ interest in STEM subjects and careers (Kier et al., 2014).

For all scales used in our survey we replaced all occurrences of “woman” and “man” with “girl” and “boy” to be more age appropriate for high school student respondents. We also added an option to select “I don’t understand the question” for each item, and a write-in space after each scale for students to share more information about why they answered the way they did or state anything they didn’t understand. Participants also completed optional demographic questions on race, gender, and LGBTQIA+ identity. Our survey design used best practices for ethical and inclusive demographic questions, including a two-part question on gender identity that allowed students to describe their current gender and sex assigned at birth, presenting demographics questions after the rest of the survey, and giving ample write-in opportunities (Jones, 2019; Tate et al., 2013). As Coburn et al. (2025) have noted, language around identity can change very quickly and demographics questions should continually evolve to reflect this. In instances where the authors perceived a lack of consensus about current terminology, we included multiple terms in the question, i.e our question included both the terms “intersex” and “difference of sexual development”.

Students who chose to participate in interviews attended an online virtual meeting with one of the researchers (CB). The semi-structured interview included questions that asked them to define biological sex in their own words, offer perspectives on the climate of LGBTQIA+ inclusion at their schools, and asked about their experience participating in the study, such as anything they did not understand in the activities or survey, or why they may have changed their answers between the pre- and post-survey (see supplemental materials for survey and interview guide).

### Student populations

We collected 318 surveys from students through classroom-based implementation working with high school teachers at three partner schools, and through online participation from students who engaged with the biology activities in a self-guided format. Two additional schools initially completed presurveys and implemented the curriculum materials but did not complete any postsurveys and thus were not included in this study. Across the three participating classrooms an average of 41.1% of the students who participated in the activities had signed parental consent forms, were present at school during all survey and implementation days, and assented to complete surveys. We conducted online participation through the Prolific platform, using the Prolific screening tools to limit our possible sample to current students, fluent in English, who were exactly 18 years old, and lived in the United States, United Kingdom, or Costa Rica (areas where we also had teacher partners). Participants who spent time outside of their regular school day to participate were offered monetary incentives through the Prolific platform. Most participants resided in the United States, across 29 different states.

The 318 surveys we collected included 93 surveys from classroom-based participants and 225 surveys from online participants. We eliminated surveys that had substantial missing data from one or more of the scales or did not have a paired pre and post occurrence, but retained surveys in which only a few questions were skipped. This resulted in an ultimate sample size of 127 completed pre and post survey pairs from 15 classroom-based participants and 112 online participants. Ten students, two from classroom-based implementations and eight from the online group, agreed to participate in interviews. In the fall of 2025, we also offered a follow-up survey to all online participants four to six months after their initial participation. There were 45 online participants who participated in this follow-up survey. Student demographic characteristics are provided in Table 2.

**Table 2a.** Sociodemographic characteristics of the student study population.

|  | LGBTQ+ |  | LGBTQ+ family |  | Transgender |  | Intersex |  |
| --- | --- | --- | --- | --- | --- | --- | --- | --- |
|  | *n* | % | *n* | % | *n* | % | *n* | % |
| Yes | 23 | 18 | 59 | 47 | 8 | 6 | 0 | 0 |
| No | 99 | 78 | 59 | 47 | 114 | 90 | 118 | 93 |
| Maybe | 5 | 4 | 7 | 6 | 2 | 4 | 3 | 2 |
| **Gender** | *n* |  |  |  |  |  | % |  |
| Girl/woman | 70 |  |  |  |  |  | 55 |  |
| Boy/man | 51 |  |  |  |  |  | 40 |  |
| Nonbinary | 4 |  |  |  |  |  | 3 |  |
| **Sex** |  |  |  |  |  |  |  |  |
| Female | 74 |  |  |  |  |  | 59 |  |
| Male | 52 |  |  |  |  |  | 41 |  |
| **Race** |  |  |  |  |  |  |  |  |
| White | 68 |  |  |  |  |  | 53 |  |
| Asian | 18 |  |  |  |  |  | 14 |  |
| Black/African American | 14 |  |  |  |  |  | 11 |  |
| Latinx/Hispanic | 9 |  |  |  |  |  | 7 |  |
| Multiple | 18 |  |  |  |  |  | 14 |  |

**Table 2b.** Sociodemographic characteristics of subsample of students interviewed.

|  | LGBTQ+ |  | LGBTQ+ family |  | Transgender |  | Intersex |  |
| --- | --- | --- | --- | --- | --- | --- | --- | --- |
|  | *n* | % | *n* | % | *n* | % | *n* | % |
| Yes | 2 | 20 | 4 | 40 | 0 | 0 | 0 | 0 |
| No | 8 | 80 | 6 | 60 | 9 | 90 | 9 | 90 |
| Maybe | 0 | 0 | 0 | 0 | 1 | 10 | 1 | 10 |
| **Gender** | *n* |  |  |  |  |  | % |  |
| Girl/woman | 7 |  |  |  |  |  | 70 |  |
| Boy/man | 3 |  |  |  |  |  | 30 |  |
| Nonbinary | 0 |  |  |  |  |  | 0 |  |
| **Sex** |  |  |  |  |  |  |  |  |
| Female | 7 |  |  |  |  |  | 70 |  |
| Male | 3 |  |  |  |  |  | 30 |  |
| **Race** |  |  |  |  |  |  |  |  |
| White | 4 |  |  |  |  |  | 40 |  |
| Asian | 3 |  |  |  |  |  | 30 |  |
| Black/African American | 2 |  |  |  |  |  | 20 |  |
| Latinx/Hispanic | 1 |  |  |  |  |  | 10 |  |
| Multiple | 0 |  |  |  |  |  | 0 |  |

### Analysis

For our quantitative analyses we used JMP student edition 19.01. For quantitative analyses of the survey data we treated answers of “I don’t understand the question” as missing data. We used confirmatory factor analysis and also calculated Cronbach’s alpha to ensure that our instruments were reliable and appropriate for our sample population (Knekta et al., 2019). A confirmatory factory analysis showed poor model fit indices for the STEM interest scale and multiple questions that did not meet the minimum recommended and factor loadings, suggesting this was not a valid instrument for our population and prompting us to remove this scale from further analysis. The four other scales showed acceptable fit indices and factor loadings, and the full results of the confirmatory factory analyses are provided in the Supplemental Materials.

We calculated a participant’s scale score by taking the mean of their Likert responses to each set of survey questions taken from the four scales. We used these four scale scores as our response variables and used generalized linear mixed models for analysis to allow for nonnormal distributions in our data. We set pre/post as a fixed effect and set participant ID as a random effect nested within context (classroom or online) and within teacher. To look further at individual level changes, we also calculated each participant’s change in score by subtracting their presurvey score from their postsurvey score for each of the five scales. These change in score values were used to create figures, and we performed Wilcoxon Signed Rank tests to determine if the mean change in score was different from zero. We also checked for significant impacts of demographics on quantitative surveys scores using Kruskal-Wallis rank sums tests.

For qualitative analysis of student interviews in Atlas.ti desktop, we first converted recordings of Microsoft Teams meetings into .mp3 audio files and linked these audio files in Atlas.ti with the autogenerated .vtt transcript files downloaded from Teams. We assigned each interviewee a pseudonym. The research team manually edited the automated transcript in Atlas.ti while listening to the audio to ensure accuracy.

We began with open coding of each interview by 2 observers, guided by our research questions and open to any emerging or unexpected insights (Saldaña, 2021). We used an open coding approach, allowing codes to emerge as we worked rather than using an *a priori* list of codes, because we wanted to remain grounded in and driven by the data during our analysis process, and we focused our process on frequent group discussion by the research team during both initial coding and a second cycle of revising and grouping codes (Saldaña, 2021). Researchers used both semantic codes, such as coding any time an interview discussed biological sex, and latent codes that pertained to our research questions such as marking statements we interpreted as expressing binarism (see Supplemental Materials for resulting codebook). During qualitative analysis, the research team met weekly to discuss any difficulties and incipient findings. We used these meetings to iteratively revise codes as we examined additional interviews, and later also compared codes between observers. It was rare for codes applied by different observers to be directly contradictory, but we did find that the diverse disciplinary backgrounds and unique perspectives of each observer led to many differences in what each coder found most significant. Occasionally codes were discarded during this iterative process as our understanding of the data deepened and if they no longer seemed relevant. Following this first round of iterative coding, (CB) created one analysis document with all observers’ codes and reviewed and merged similar codes, added additional code groupings and subcodes, and identified interrelationships and patterns among codes and participant demographics. Following this, all researchers then examined the updated analysis document together, along with all observers’ research memos, made additional edits to coding and groupings as needed, and reached a synthesized consensus of the key findings through group discussion.

Our qualitative and quantitative analyses were conducted over the course of one year and our approach as a research team was to let emerging results from each methodological approach inform the process of the other. For example, we conducted quantitative analyses of preliminary data in Summer of 2024 and our research team had initial trends and lingering questions in mind as we revised and implemented the interview guide. During qualitative analysis of interviews, the research team began to note the trend of the varied ways interviewees discussed and defined sex and gender and this encouraged us to fully explore differences among students in quantitative analysis of presurveys. We present our findings of qualitative and quantitative analyses together because this reflects our simultaneous mixed-methods approach.

## FINDINGS

We collected data from 127 high school students who submitted both complete pre- and postsurveys from before and after their participation in the inclusive biology activities. We checked the reliability of our four scales using Cronbach’s’ alpha test. The Heteronormative beliefs: normative behavior scale (α=0.84), the Heteronormative beliefs: essential, binary sex and gender scale (α=0.92), the Transphobia scale (α=0.93), and the Gender /Sex diversity affirmation scale (α=0.97) all showed high reliability in our study. Although it did not affect overall reliability, we did note that 13% of surveys had missing data or a response of “I don’t understand the question” for question six on the transphobia scale: “I believe that the male/female dichotomy is natural.” Several respondents used write-in spaces to indicate that they did not know what the word dichotomy meant, and we recommend that the wording of this question should be changed in future research.

### RQ1. Most students’ attitudes and beliefs changed after the curriculum intervention

We found that there was a significant difference in survey scores from pre to post for the Heteronormative beliefs: essential, binary sex and gender scale (*F*_1,126_ = 24.6, *p* < 0.0001). There was also a significant difference in survey scores from pre to post for the Gender/sex diversity affirmation scale (*F*_1,125.2_ = 21.1, *p* < 0.0001). Parameter estimates for all factors are in Table 3. Looking at individual participant’s changes in scores from pre to post, we saw that participants had a mean change of -0.41 ± 0.08s.e. on the 7-point Likert scale from pre to post for the Heteronormative beliefs: essential, binary sex and gender scale, and this change in scores was significantly different from zero (*W* = -1850.0, *p* < 0.0001, Figure 2). Participants showed an increase in attitudes of affirmation of sex and gender diversity significantly different from zero *W* = 2095.5, *p* < 0.0001, with a mean change of 0.33 ± 0.07s.e. on the 7-point Likert scale from pre to post (Figure 2). We found no significant statistical difference in scale scores for the Transphobia and Heteronormative beliefs: normative behavior scales between pre- and postsurveys. For students who completed the follow-up survey, their survey scores remained significantly different from their presurvey scores, indicating that the shifts in attitudes and beliefs were durable over time several months following the curriculum intervention (Figure 3, Table 4). Pre-survey scores differed from post-survey scores, and post-survey scores were not significantly different from follow-up scores (Table 4).

**FIGURE 2.**
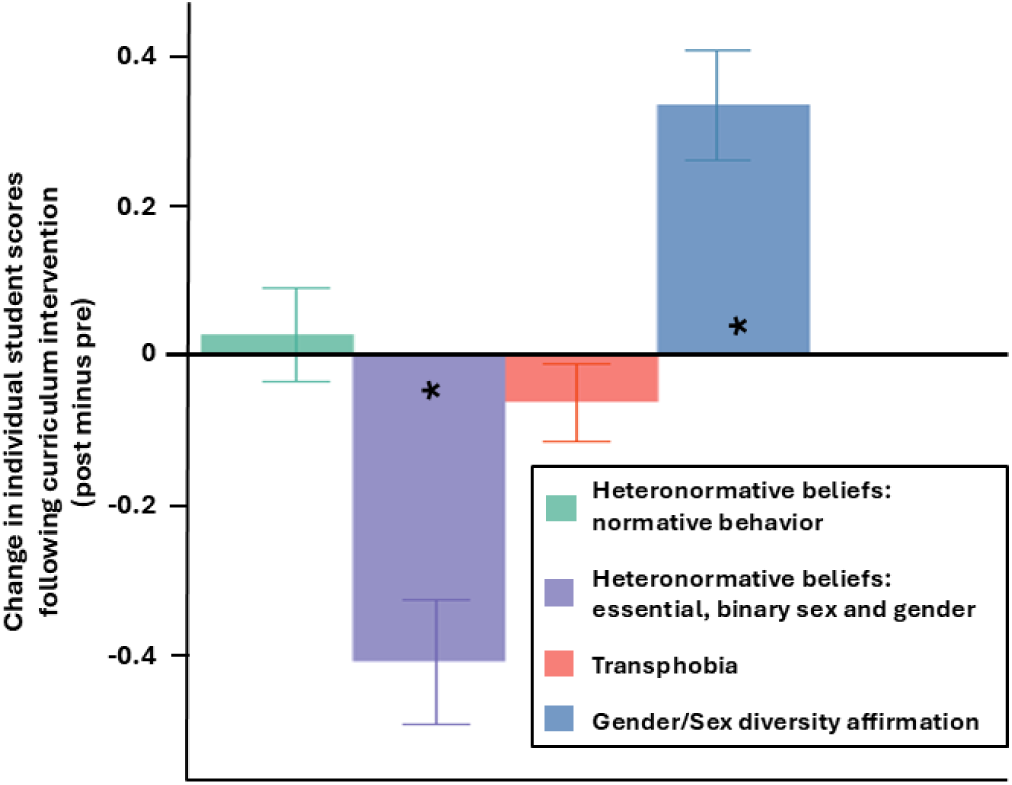
Changes in individual student scores for Heteronormative beliefs: essential, binary sex and gender scale, Heteronormative beliefs: normative behavior scale, Transphobia scale, and Gender/sex diversity affirmation scale following participation in inclusive biology activities, calculated as post minus pre survey scale scores with bars showing standard errors. Asterisks denote significant difference from zero (Wilcoxon signed rank test). Note that the Transphobia scale was measured on a 6- point Likert scale, while the other variables were measured on a 7-point Likert scale.

**FIGURE 3.**
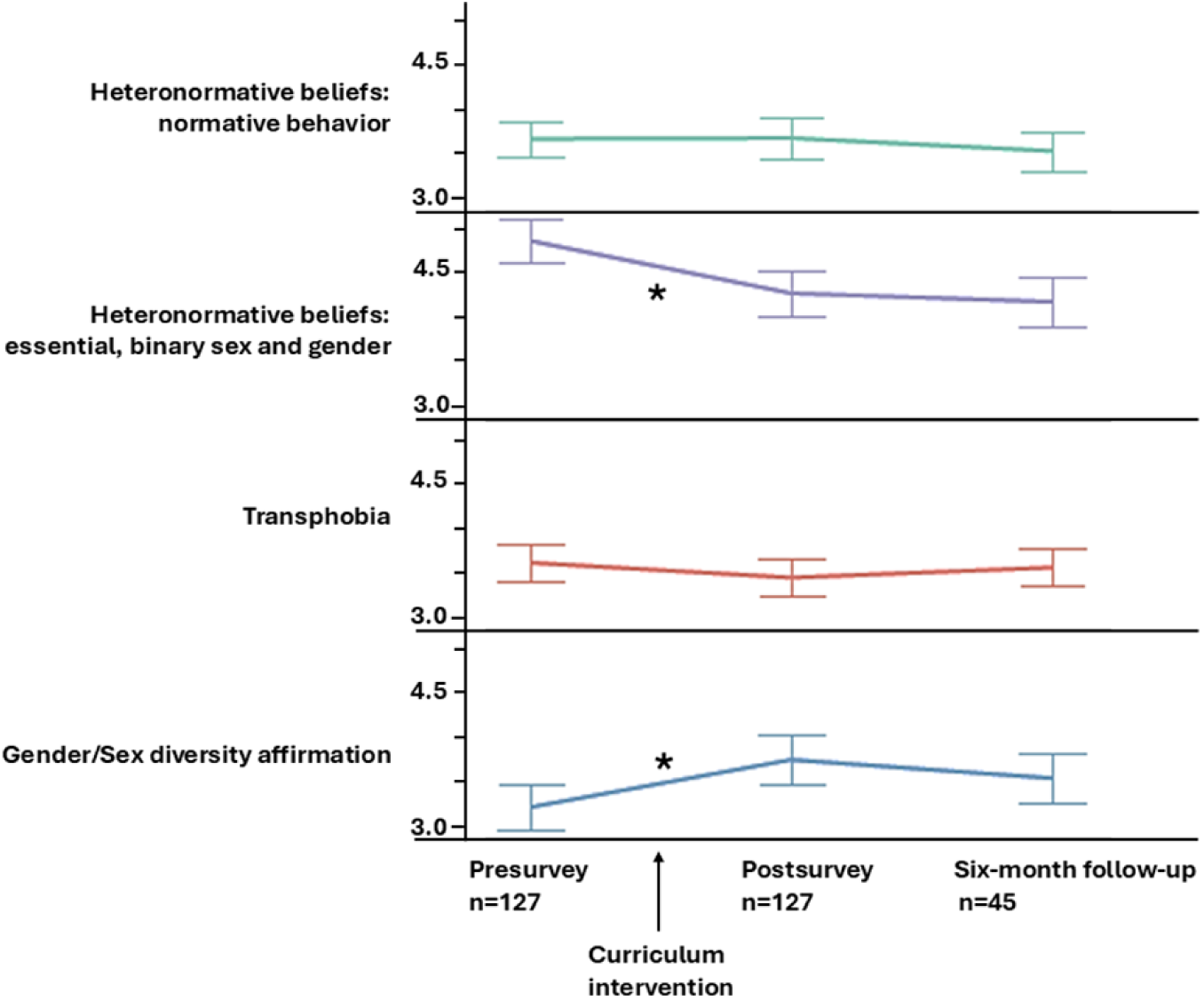
Means and standard errors for the Heteronormative beliefs: essential, binary sex and gender scale, Heteronormative beliefs: normative behavior scale, Transphobia scale, and Gender/sex diversity affirmation scale over time, from presurvey, postsurvey, and a follow-up survey four to six months following the curriculum intervention. Asterisks denote significant difference between time points. Note that the Transphobia scale was measured on a 6-point Likert scale, while the other variables were measured on a 7-point Likert scale.

**Table 3.**
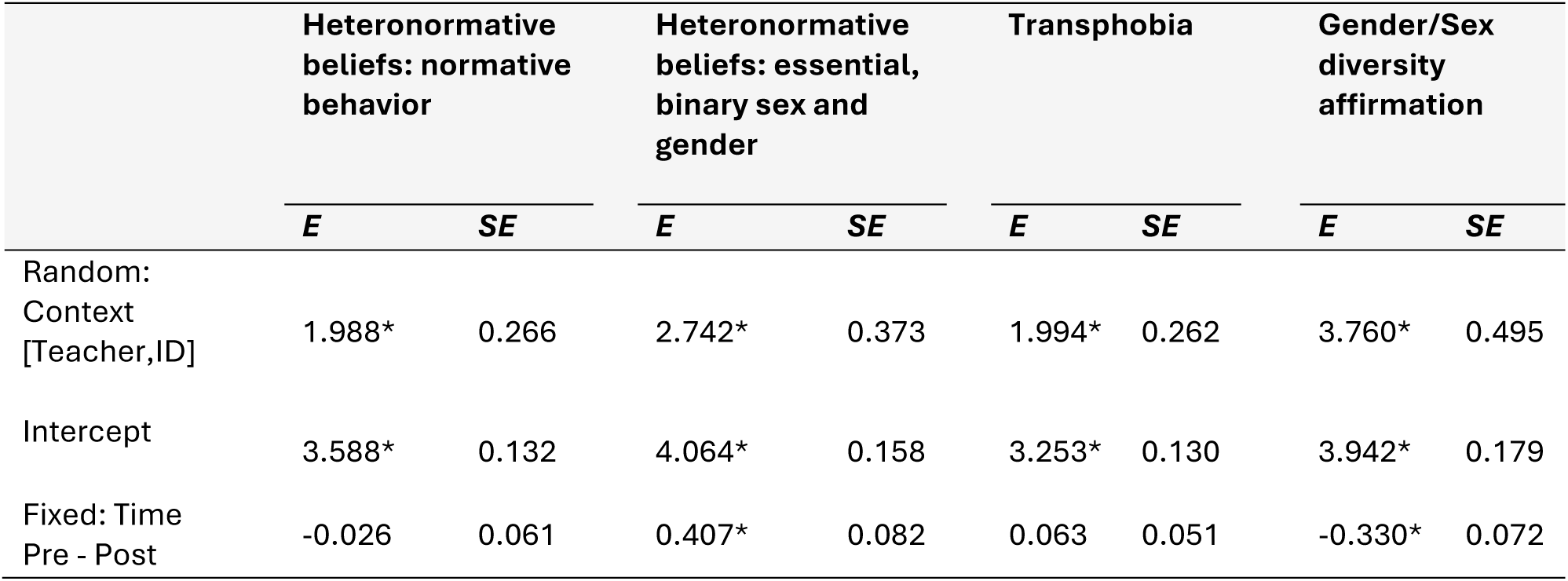
Effects on student attitudes and beliefs on four scales as measured by pre- and postintervention surveys, analyzed with GLMM. The model includes a random effect of student ID and teacher nested within learning context (classroom or online), and the fixed effect of pre or postintervention. Parameter estimates (*E*) and standard errors (*SE*) are listed for each effect with significance denoted by * (p<0.0001). N=127

|  | Heteronormative beliefs: normative behavior |  | Heteronormative beliefs: essential, binary sex and gender |  | Transphobia |  | Gender/Sex diversity affirmation |  |
| --- | --- | --- | --- | --- | --- | --- | --- | --- |
|  | *E* | *SE* | *E* | *SE* | *E* | *SE* | *E* | *SE* |
| Random: Context [Teacher, ID] | 1.988* | 0.266 | 2.742* | 0.373 | 1.994* | 0.262 | 3.760* | 0.495 |
| Intercept | 3.588* | 0.132 | 4.064* | 0.158 | 3.253* | 0.130 | 3.942* | 0.179 |
| Fixed: Time Pre - Post | -0.026 | 0.061 | 0.407* | 0.082 | 0.063 | 0.051 | -0.330* | 0.072 |

**Table 4.** Effects on student attitudes and beliefs on four scales as measured by surveys at three time points: pre- and post intervention, and a follow up four to six months later. The GLMM model includes a random effect of student ID and the fixed effects of survey time. Parameter estimates (*E*) and standard errors (*SE*) are listed for each effect with significance denoted by * (p<0.0001). N=45

|  | Heteronormative beliefs: normative behavior |  | Heteronormative beliefs: essential, binary sex and gender |  | Transphobia |  | Gender/Sex diversity affirmation |  |
| --- | --- | --- | --- | --- | --- | --- | --- | --- |
|  | *E* | *SE* | *E* | *SE* | *E* | *SE* | *E* | *SE* |
| Random: ID | 1.987* | 0.440 | 2.455* | 0.564 | 1.811* | 0.408 | 2.745 | 0.640 |
| Intercept | 3.633* | 0.220 | 4.860* | 0.257 | 3.569* | 0.217 | 3.160 | 0.278 |
| Fixed: Time Post-Pre | 0.009 | 0.095 | -0.563* | 0.153 | -0.173 | 0.117 | 0.549* | 0.183 |
| Fixed: Time Follow up-Post | 0.033 | 0.101 | 0.056 | 0.158 | 0.118 | 0.117 | -0.213 | 0.182 |

In interviews, the majority of students expressed that they learned things in the lessons they never knew before or that the lessons expanded their understanding. Several students also had positive feedback for the impact of the lessons overall, saying they found them interesting and felt they would retain the information better than some of the “boring” assignments from their other past science courses. As we considered changes in students’ attitudes, we looked at qualitative and quantitative responses together for individual students, especially those who did not fit the dominant trend of increasing affirming attitudes. We generally found cohesive agreement of individual attitudes and beliefs expressed in surveys and with sentiments expressed in interviews; i.e. students who expressed strongly binary views of sex in their interviews also scored high on the Heteronormative beliefs: essential, binary sex and gender scale. However, students were less consistent between surveys and interviews in discussing whether there had been any change in their thinking in response to the lessons. Some students who reported in interviews that their thinking did not change following the activities actually showed large quantitative shifts in their survey scores. For example, for the four students who claimed in interviews their attitudes and beliefs had not changed following the curriculum intervention, the mean change in their scores on the Heteronormative beliefs: essential, binary sex and gender scale was -0.633 ± 0.41s.e. This contradiction coincides with a complex feeling that several interviewees expressed, which seemed to indicate they were struggling to reconcile information they had been presented with in the lessons with their previous beliefs. For example, Mei stated, “Like my perspective kind of grew, if that makes sense. But my [survey] answers pretty much stayed the same…” In parallel, Abbas stated in his interview that, “My opinion before what I read was completely different to what my opinions afterwards…my thinking changed quite a lot.” But Abbas also maintained in his interview that he firmly believed there were only two genders and volunteered the unsolicited assertion that he thought people could not change genders. To further highlight his conflicted feelings, Abbas’s quantitative survey scores actually changed the least of any of the interviewees.

### Humans and nonhumans

We also saw that several interviewees attempted to reconcile their internal conflict about whether, and how, their attitudes might be shifting following the curriculum intervention by demarking a distinct separation between humans and animals. Abbas seemed to be thinking aloud during his interview, musing, “In nature, there might be more than two genders. I mean, humans are also part of nature, but I mean in *raw* nature it might be different [than in humans].” It is interesting to note that several interviewees focused on the biology of humans in their answers even though the activities we presented to them were mostly focused on animals and none of our interview questions used the word gender or asked about the biology of sex specifically in humans. The tension between humans and nonhumans also came up when we asked Ruby about whether she thought our biological activities would be a welcome addition to the curriculum in her school environment:

I think when it comes to animals being a sex that’s not just male and female, I think people are more willing to open their minds to that. So I don’t think that would be, like controversial. But I think people find it more… controversial or different to their views when that same mindset is applied to humans.

Although it was not an intended part of our data collection, we also noted that several teachers who requested to use our materials but did not participate in data collection were more confident about their ability to implement activities that did not contain any content about humans.

#### RQ2. Straight and LGBTQIA+ students differed in their attitudes and beliefs

We were also interested in understanding how students’ demographics might impact their attitudes and beliefs about the biology of sex, both before and after exposure to the curriculum intervention. Thus, we checked for significant impacts of demographics in both the initial presurvey measurements of students’ understanding of the biology of sex and in postsurveys and interviews. We found that having an LGBTQIA+ family member or close friend influenced participant presurvey scores compared to those who did not have an LGBTQIA+ family member for the Heteronormative beliefs: essential, binary sex and gender scale (χ^2^ = 21.5, *p* < 0.0001), Heteronormative beliefs: normative behavior scale, (χ^2^ = 29.8, *p* < 0.0001), Gender/sex diversity affirmation scale (χ^2^ = 27.4, *p* < 0.0001), and Transphobia scale (χ^2^ = 26.6, *p* < 0.0001). Survey participants who had LGBTQIA+ family members came in with lower binary sex and gender essentialist attitudes and more diversity-affirming attitudes, as evidenced by their pre survey scores (Figure 4). Similarly, students who stated that they were LGBTQIA+ themselves had significantly lower scores on the Heteronormative beliefs: essential, binary sex and gender scale (χ^2^ = 25.4, p < 0.0001), Heteronormative beliefs: normative behavior scale, (χ^2^ = 18.2, *p* = 0.0001), and Transphobia scale (χ^2^ = 20.7, *p* < 0.0001), and higher scores on the Gender/sex diversity affirmation scale (χ^2^ = 24.8, *p* < 0.0001) in their presurvey responses (Figure 5). Aside from the distinct patterns for LGBTQIA+ individuals and participants with LGBTQIA+ family or close friends, we saw no significant influence of any other recorded participant demographic characteristics (e.g. race, sex, gender).

**FIGURE 4.**
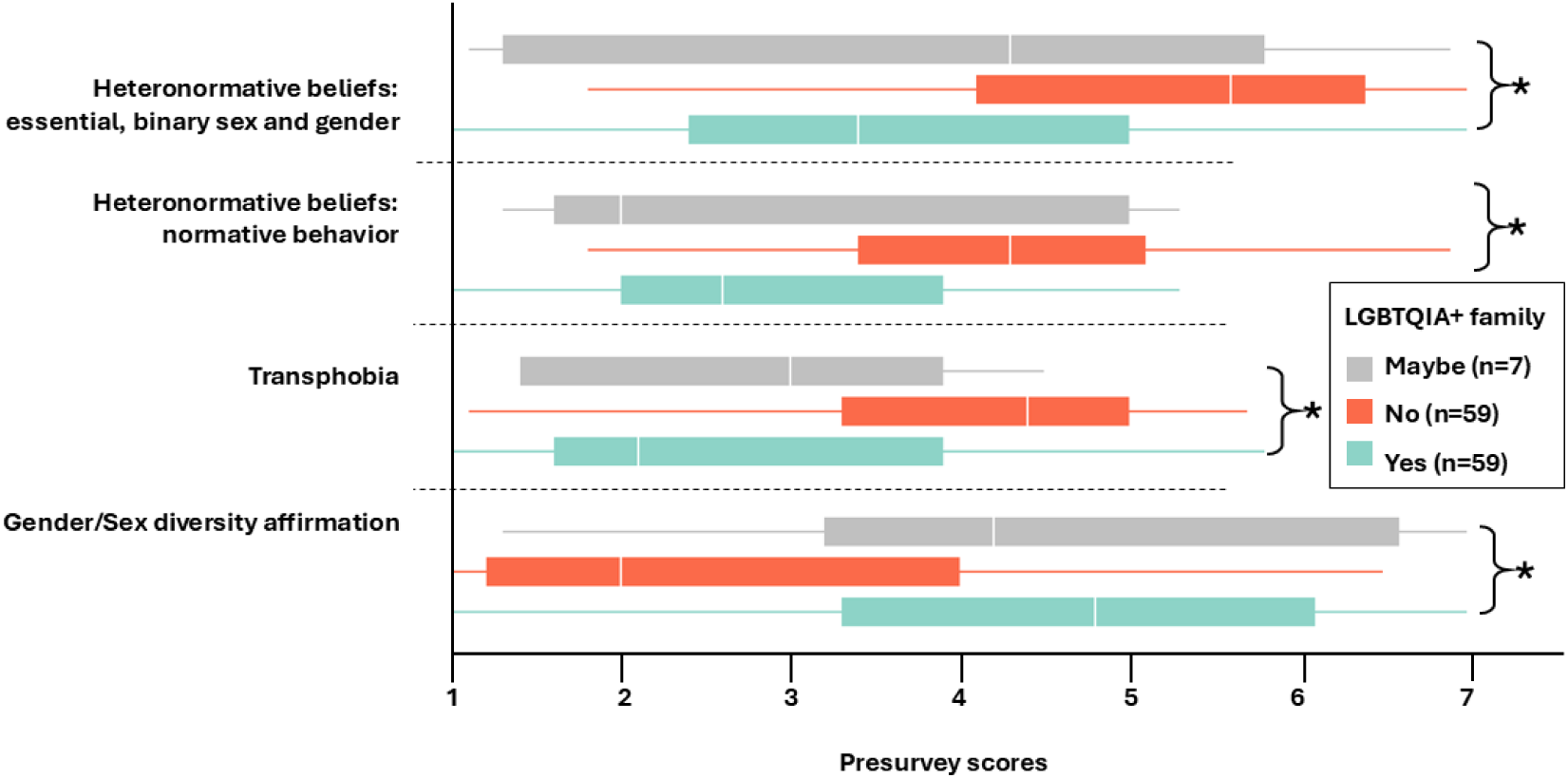
Box plot showing that presurvey scores differed for students with and without LGBTQIA+ family member for the Heteronormative beliefs: essential, binary sex and gender scale, Heteronormative beliefs: normative behavior scale, Transphobia scale, and Gender/sex diversity affirmation scale. White line indicates the median and asterisks denote significant difference between groups. Note that transphobia was measured on a 6-point Likert scale, while the other variables were measured on a 7-point Likert scale.

**FIGURE 5.**
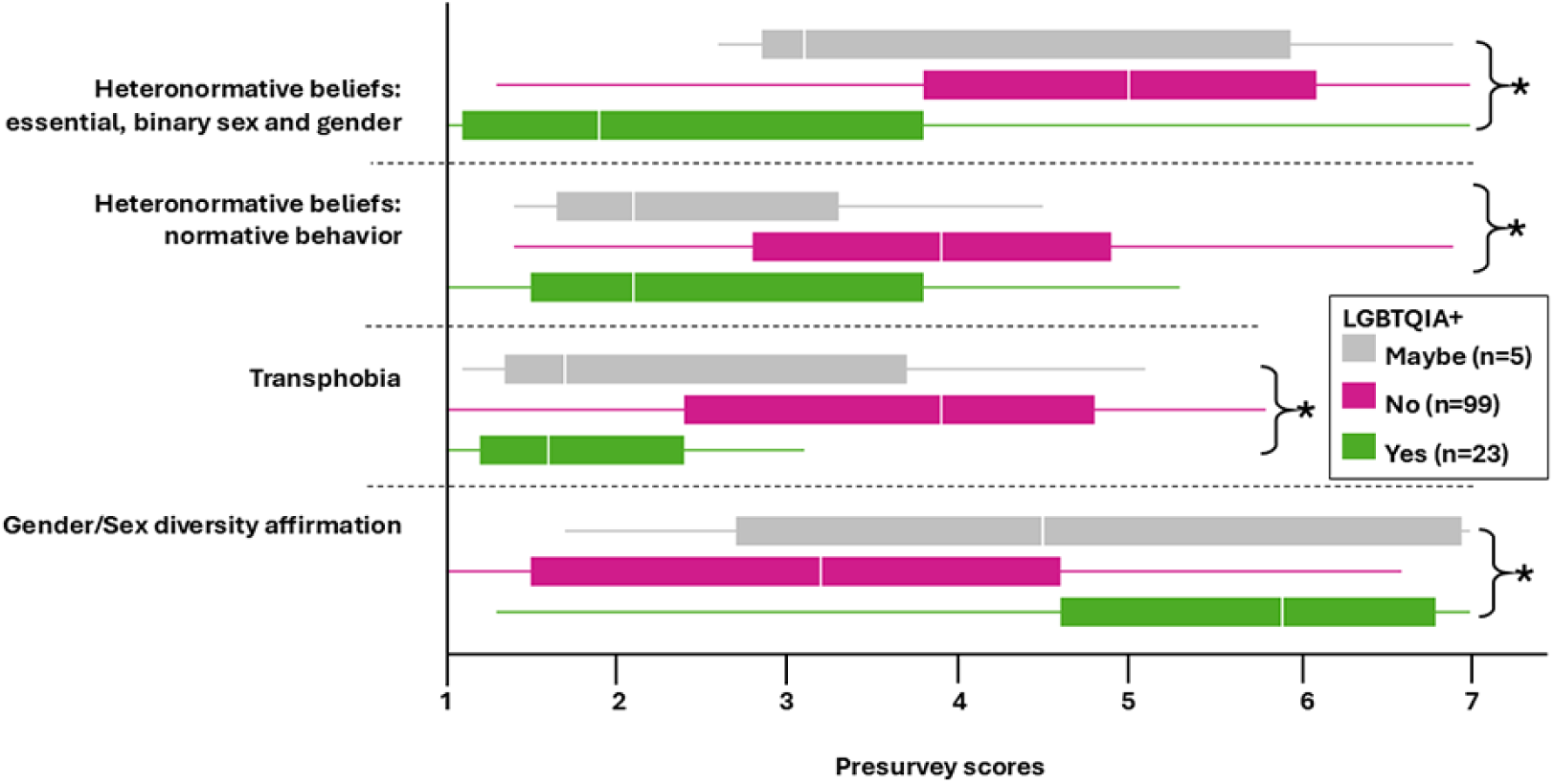
Box plot showing that presurvey scores differed for straight and LGBTQIA+ students for the Heteronormative beliefs: essential, binary sex and gender scale, Heteronormative beliefs: normative behavior scale, Transphobia scale, and Gender/sex diversity affirmation scale. White line indicates the median and asterisks denote significant difference between groups. Note that transphobia was measured on a 6-point Likert scale, while the other variables were measured on a 7-point Likert scale.

Consistent with the distinct quantitative findings for LGBTQIA+ individuals, we saw marked differences between students who described themselves as straight and those who described themselves as LGBTQIA+ in the way they discussed the biology of sex in interviews. When speaking about the biology of sex, especially in the way they described their understanding of male and female, the two LGBTQIA+ students we interviewed focused on the type of sex cells produced and the role of the individual in procreation. For example, Julia explained:

Males produce cells that go into the female to help. That’s how I see it, to classify it. So if there is something, an organism, that provides those cells, I see it as male. And if it can change itself into the female type, then I then see it as female.”

Some of both the straight and LGBTQIA+ students mentioned sex chromosomes and also talked about sex as a quality that an individual is born as or starts with. However, LGBTQIA+ interviewees acknowledged that sex chromosomes can also come in atypical combinations and that an individual’s sex can potentially change over time, at least for some organisms. Only straight students mentioned sex hormones, external genitalia, and innate behavioral patterns in discussing their definitions of biological sex during interviews, as in the response below from Aliya when asked how she would you define biological sex in her own words:

Umm, I think it’s the gender you were born. I would say it’s maybe the gender you are biologically born, like you’re born male or female. It’s down to your innate characteristics, your innate…not characteristics… your innate behaviors I think.

Some straight students also conflated sex and gender in their interpretation of, and response to, the interviewer prompting them to define biological sex, as also demonstrated by Aliya’s response above. This difference between LGBTQIA+ students and their straight peers is also apparent in responses to a particular item on the Heteronormative beliefs: essential, binary sex and gender scale: “Gender is the same thing as sex.” Responses to this survey item varied significantly in presurveys (χ^2^=16.2, p=0.0003), with straight students much more likely to agree with this statement than LGBTQIA+ students.

#### RQ2. Straight and LGBTQIA+ students have different experiences in science education

We also explored how students’ LGBTQIA+ identity impacted the way students responded to the curriculum intervention and what factors may relate to different experiences of the curriculum intervention and in science education across identity. We found that there was a significant interaction between survey time from pre to post and LGBTQIA+ identity that impacted student scores (Table 5). For most variables, changes in scores following the curriculum intervention were in the same direction across LGBTQIA+ identity (Figure 6). For the Heteronormative beliefs: normative behavior scale, straight students showed a trend towards a small increase in heteronormative beliefs, while LGBTQ students showed a trend towards a decrease in heteronormative beliefs, however, the mean changes in this scale were not significantly different from zero for any of these groups (LGBTQIA-Maybe: *W* = -3.00, *p* = 0.500, LGBTQIA-No: *W* = 417.00, *p* = 0.146, LGBTQIA-Yes: *W* = -46.00, *p* = 0.163). Straight students also tended to have larger decreases in their scores on the Heteronormative beliefs: essential, binary sex and gender scale, with a mean change of -0.389 ± 0.09s.e., as compared to LGBTQIA+ students who had a mean change in scores of -0.252 ± 0.159s.e. For students who were LGBTQIA+, this reduction in heteronormative, essential, binary sex and gender beliefs was not statistically significant from zero, driven in part by this groups’ very low initial scores on this scale (Figure 6). For example, over half of LGBTQ+ students had a mean score between 1 and 2 on the Heteronormative beliefs: essential, binary sex and gender scale before the intervention, meaning there was no room on the scale for their scores to decrease any further. We hesitate to draw any conclusions for students who are answered “Maybe” about their LGBTQIA+ identity because of the small sample size (N=5).

**FIGURE 6.**
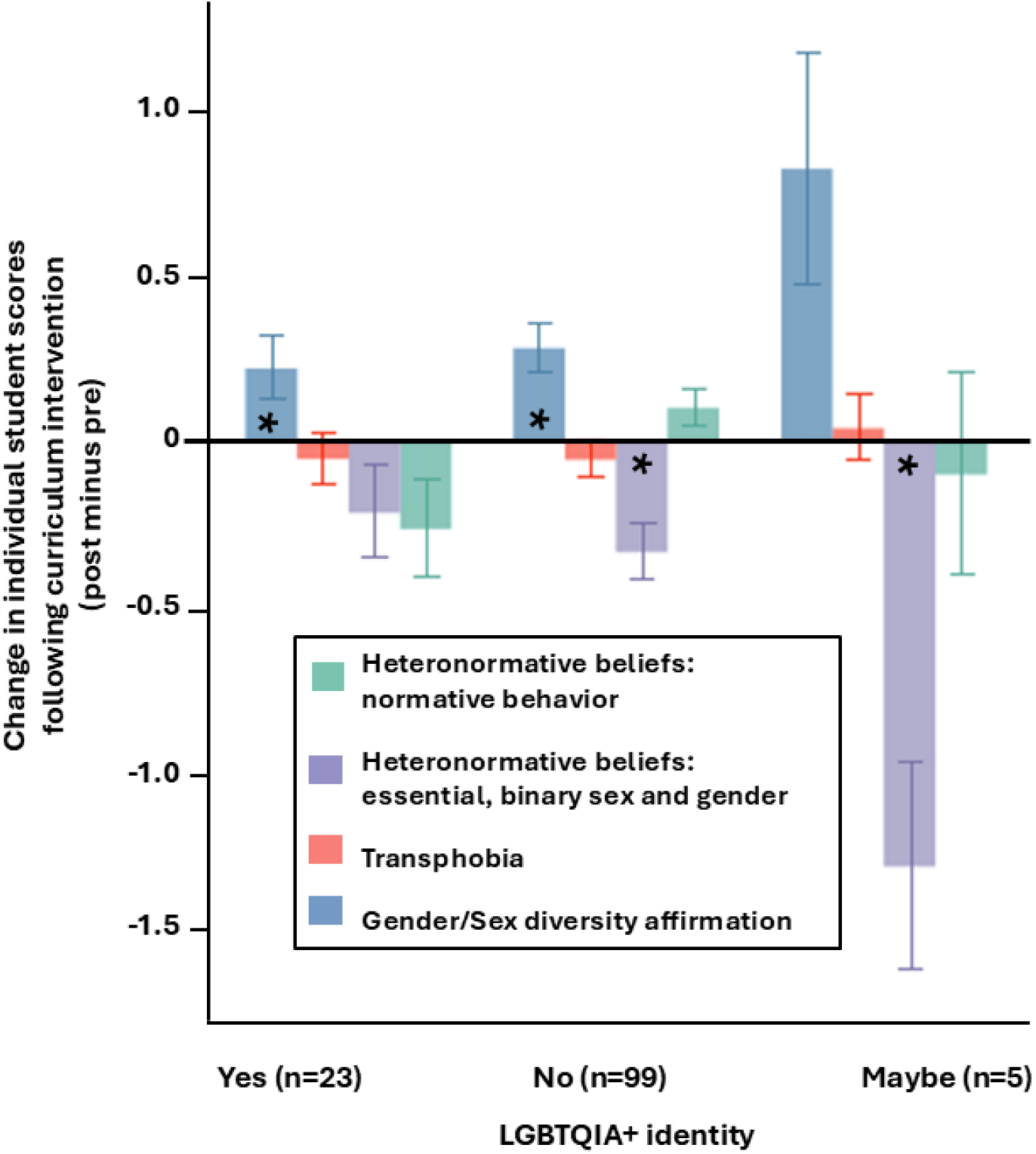
Changes in individual student scores following participation in inclusive biology activities, calculated as post minus pre survey scale scores with bars showing standard errors, separated by LGBTQIA+ identity of the students. Asterisks denote significant difference from zero (Wilcoxon signed rank test). Note that the Transphobia scale was measured on a 6-point Likert scale, while the other variables were measured on a 7-point Likert scale.

**Table 5.** Effects on student attitudes and beliefs as measured by surveys pre- and post intervention, with interaction of student LGBTQIA+ identity. The model includes a random effect of student ID and teacher nested within learning context (classroom or online), and the fixed effects of survey time, LGBTQIA+ identity, and their interaction. Parameter estimates (*E*) and standard errors (*SE*) are listed for each effect with significance denoted by * (p<0.05). N=128

|  | Heteronormative beliefs: normative behavior |  | Heteronormative beliefs: essential, binary sex and gender |  | Transphobia |  | Gender/Sex diversity affirmation |  |
| --- | --- | --- | --- | --- | --- | --- | --- | --- |
|  | *E* | *SE* | *E* | *SE* | *E* | *SE* | *E* | *SE* |
| Random:<br>Context [Teacher, ID] | 1.623* | 0.221 | 2.062* | 0.289 | 1.630* | 0.218 | 2.991* | 0.401 |
| Intercept | 2.252* | 0.284 | 2.422* | 0.328 | 2.030* | 0.280 | 5.648* | 0.379 |
| Fixed: Time<br>Pre-Post | 0.309* | 0.141 | 0.252 | 0.188 | 0.065 | 0.121 | -0.248 | 0.167 |
| Fixed: LGBTQIA<br>Maybe-Yes | 0.028 | 0.671 | 0.283 | 0.776 | 0.340 | 0.662 | 0.052 | 0.898 |
| Fixed: LGBTQIA<br>No-Yes | 1.712* | 0.315 | 2.095* | 0.364 | 1.549* | 0.310 | -2.191* | 0.396 |
| Fixed: Time * LGBTQIA<br>Pre-Post*Maybe-Yes | -0.188 | 0.333 | 1.228* | 0.446 | -0.105 | 0.288 | -0.692 | 0.396 |
| Fixed: Time * LGBTQIA<br>Pre-Post*No-Yes | -0.420* | 0.156 | 0.137* | 0.209 | 0.002 | 0.135 | -0.070 | 0.186 |

Our interviews also uncovered nuance in how students’ sociocultural identities may impact their experiences in STEM education, their response to the curriculum intervention, and how previous experiences in science education environments may relate to their attitudes and beliefs around sex and gender. In talking about previous experiences in their science education, many interviewees noted that some of the content of our activities was new or different than what they had learned in previous science classes, although they had covered similar topics before. Several interviewees mentioned learning about some of the material in our lessons before, through online sources or other sources outside of school. One student, a Latinx 12^th^ grader who identified as LGBTQIA+, spoke explicitly about the impact of her science instructors’ use, or not, of inclusive teaching approaches:

Charlie (interviewer): Do you think that queer people or LGBTQIA+ people feel comfortable in science class? Like, do you think there’s anything that makes LGBTQIA+ people more or less likely to go into science?

Ruby: I think in the classes that I’ve been, I can see why it would be harder, difficult. Because a lot of the way that, at least when it comes to sex and gender, that is explained, it’s very black and white. Because this aligns, you know, with the typical way we think of nature, and that there’s only male and female. Because the main way that they talk about biology is through having two sexes and then the way those reproduce. I think it can be hard because that puts things, like, it makes it in a heteronormative lens. Which I think can be…it can make queer students feel left out or kind of like they’re not part of the…

When *I know* that in nature it’s not just male and female relationships. Same sex relationships are common in nature. So, I think that the fact that that isn’t included can make queer students feel left out and things like that. So I think that can be hard, but there’s also been classes I’ve taken that have been more inclusive of that. So I think it depends. But I think kind of normalizing the fact that there’s different relationships that happen in nature or there’s different identities that can be seen in nature, I think that can make it easier for people to feel included and not feel, like, out of the norm. Because they are common occurrences, and I think it’d be better overall to include that more in schools.

Charlie (interviewer): Did you say that you’ve had some classes where they did talk more about that? Like, more inclusive than other classes?

Ruby: Yeah, it depended. Because there were some teachers that I knew that were like, more old-school. So they just stuck to the curriculum or textbook. But since those are, like, old, they don’t really include things like this that are more open to the nuances.

Compared to, I know some teachers are more aware of that and stuff, so they tried… If they were talking about sexes that occur naturally, they might go off from the curriculum and be like, "There’s male and female that are usually XY and XX," but then they themselves would be like "But that’s not the only possibilities." So, it’s not usually something that’s included. But they would go out of their way to include that, because they know that it’s better to be aware of that and to be informed of that.

Earlier in her interview, Ruby mentioned that she knew about some of the material in our curriculum intervention lessons because she had learned it on her own through online sources. Her interview responses seem to indicate that this knowledge has empowered her with the feeling that queer identities are completely natural, which makes it even more alienating when her “old-school” science teachers leave out what she knows to be true. Ruby articulated what it was like to be a well-informed queer student who is let down by an outdated science curriculum and teachers who made little effort to move beyond faulty generalizations in their teaching. Ruby demonstrated a depth of knowledge of biology in our interview that was gained through independent learning online in her free time, indicating she may have had at least some prior interest in science. Yet, in her concluding interview remarks she off-handedly mentioned, “I’m not usually the biggest fan of, like, science, and stuff like that.”

Interestingly, another student who also identified as LGBTQIA+, Julia, expressed views that contrasted with Ruby. Julia explained that she thought LGBTQIA+ students should feel comfortable in science class due to what she saw as the separation between the scientific discipline and students’ sociocultural identities, as maintained by teachers who keep “…a nice division between the material we’re learning and us as the students.” She told us that personal identities should not be consequential in science classes because, “science class is focused more on what the actual lesson is rather than separating that LGBTQIA+ person from the social area, or trying to incorporate them into that lesson somehow.” Julia’s view seems fueled by a desire to avoid a focus on her LGBTQIA+ identity that would make her feel othered or separate from the social group. The contrast between Julia and Ruby’s perspective illustrates that there is not a universal consensus of opinion within the LGBTQIA+ community when it comes to the best inclusive teaching strategies in science education, and shows that some LGBTQIA+ students might find safety in an approach to science education that ignores sociocultural context because it keeps their identity invisible and out of the way of their education. Although they favor different solutions, both Julia and Ruby’s perspectives show that anti-LGBTQIA bias and/or the anticipated threat and avoidance of it has affected both their science educational experiences.

On the other hand, nearly all of the straight interviewees stated that LGBTQIA+ students are welcomed and should feel comfortable in science class, and saw no reason why LGBTQIA+ students should be any more or less likely to pursue science in their education or career than straight students. One straight interviewee affirmed that he thought his science classes are welcoming: “I only know of one guy and everyone was pretty calm with him. Like no one really cared if he was gay or not.” However, this interviewee also suggested this gay person might tend towards a subject that was “more creative” rather than pursuing science. These findings suggest that some stereotypes about LGBTQIA+ students’ academic interests may persist among their peers, and that many straight students remain unaware of how their LGBTQIA+ peers may be experiencing their sciences classes differently.

The majority of interviewees thought the material in the curriculum intervention would be welcome at their school, or at least some parts would be welcome if only the sections focused solely on animals and not humans. Several students indicated awareness of the broader social and historical context in which their education takes place, and suggested that inclusive approaches to education are more accepted nowadays than in the past. One exception was Michael, who explained that he went to a Christian school where the type of lessons we showed him would not be welcome, and where any ideas other than a two-sex viewpoint determined by chromosomes would not be allowed. Michael also declared honestly, if somewhat apologetically, that “as bad as that is,” that, “generally, no”, LGBTQIA+ students are not welcomed due to the religious and “conservative” nature of his school.

To return to our research questions, our qualitative and quantitative findings support that our curriculum intervention did impact attitudes and belief of the students (RQ1). Students increased in their affirming attitudes of sex and gender diversity as well as decreased in their heteronormative, essential sex and gender beliefs (RQ1). Our qualitative analysis uncovered that this shift was likely due to the students being exposed to new information in the curriculum intervention that was different than what they had learned about these topics in the past (RQ1). In addressing our second research question, we saw interesting differences in attitudes among LGBTQIA+ students and their straight peers that were present in presurveys, before engaging with our curriculum intervention (RQ2). Our qualitative and quantitative analysis also revealed differences in the ways that LGBTQIA+ students and straight students understood and talked about the biology of sex and their science education experiences (RQ2).

## DISCUSSION

### A critical pedagogical approach is needed

Our study reaffirms that students are learning more than just science in science class, whether or not their educators chose to take notice of this hidden curriculum (Giroux & Penna, 1979; Jackson, 1990; Martin, 1976). Rather than ignore the sociocultural context in which education occurs, we assert that a critical pedagogical perspective is needed, one that recognizes education inherently involves many opportunities to either uphold or challenge the status quo, and that school can play an integral role in challenging or reinforcing normative enculturation (Butler, 1999; Freire, 2000; Keenan, 2017; Rende Mendoza & Johnson, 2024). Our study offers evidence that practical and concrete changes to biology curriculum content have the power to reduce the sway of bioessentialist fallacies and increase student attitudes of affirmation for gender and sexual diversity.

### Inclusive science educational interventions can effectively shift student attitudes

Our findings show that even a brief exposure to inclusive biology curriculum content was effective in significantly reducing students’ heteronormative, bioessentialist thinking about sex and gender. Some of the study participants spent less than one hour engaged in the inclusive biology activities, and this brief encounter still elicited significant shifts that were apparent in student survey scores and in some of their interviews, including in surveys 6 months later. This result aligns with previous work by Donovan et al. (2019) that showed even a brief reading about genetics that was carefully crafted to refute neurogenetic essentialism could reduce girls’ belief that science ability is innate, which was also associated with their level of STEM interest.

However, previous research suggests that sustained incorporation of inclusive practices throughout the design and implementation of a whole curriculum is preferable to one-off lessons that can be perceived as inauthentic and tokenizing (Long et al., 2021; Morgan et al., 2024). The length and quality of intervention is likely important in determining whether shifts in student attitudes will remain robust in the long term and how shifts in attitude may lead to reductions in discriminatory behavior. Although we saw encouraging results in our study from brief exposure to curriculum interventions, we suggest that interspersing our modified lesson plans throughout the school year along with other inclusive teaching practices would be the preferable approach.

Some studies have shown that shifts in attitudes as the result of brief interventions may not be long lasting, and that relationships between changes in implicit bias, explicit attitudes, and discriminatory behaviors are not always strongly correlated (Forscher et al., 2019; Paluck et al., 2021; Paluck & Green, 2009). For example, Wüthrich et al. (2024) implemented a curriculum-based intervention designed to reduce negative attitudes towards peers with disabilities, but saw a reduction only in students’ explicit bias without a significant reduction in implicit bias. A longer intervention designed to reduce gender bias through a 36 session curriculum over a 5 month period achieved reductions in both implicit and explicit gender bias (Lu et al., 2025). In our study, the internal conflicts described in interviews by a few of our participants whose survey results seemed to contradict their interview statements about whether or not their thinking had changed suggest that more research is needed on the relationship between student knowledge, implicit and explicit attitudes, and discriminatory behavior. Our findings from students who participated in the follow-up survey four to six months following exposure to the curriculum intervention suggest that the changes in attitudes we measured remained durable over time. We hope future research will continue to explore whether receiving new scientific information through a curriculum intervention that controverts pseudoscientific beliefs is sufficient to prevent students from returning to, or acting on, bias that was informed by their previous beliefs over the long term.

Interestingly, we found significant changes on the Heteronormative beliefs: essential, binary sex and gender scale but not on the Heteronormative beliefs: normative behavior scale, which are two subscales originally from the same source (Habarth, 2015). We also saw that there were some differences in the way LGBTQIA+ and straight students responded to these scales both before and after the curriculum intervention. Some straight students actually increased in their scores on the Heteronormative beliefs: normative behavior scale, and while these changes were small and not significantly different from zero, it could account for why we saw no significant reductions in the Heteronormative beliefs: normative behavior scale when looking at the full study population. The significant change following the intervention in the Heteronormative beliefs: essential, binary sex and gender scale was driven by a significant reduction in these attitudes by straight students, with a nonsignificant trend in the same direction for LGBTQIA+ students, many of whom who started out with very low scores on this scale to begin with. These findings suggest that there are complex factors affecting these attitudes for students, including individual sociocultural identity, and that there may be complex relationships among the different aspects of heteronormative beliefs.

We saw no significant changes in the Transphobia scale following the curriculum intervention. One possible reason for this is that the transphobia scale measures attitudes that are more explicitly about humans, such as item 8, “I believe a person can never change their gender.” In contrast, some of the other scales contain prompts that may or may not be interpreted as being specifically or exclusively about humans, including prompts about the nature of biological sex in general, such as the items “Sex is complex; in fact, there might even be more than 2 sexes” from the Heteronormative beliefs: essential, binary sex and gender scale, and “Biological sex is not just female or male; there are many possibilities” from the Gender/Sex diversity affirmation scale.

### Humans as animals and the naturalistic fallacy

Our findings point to an important area for further exploration in how students understand humans within the natural world. Much of the content of the inclusive biology lessons was about nonhuman organisms, although some students did participate in activities that included some content about human biology, and our survey questions did prompt students to think about their views within a human social context. One of the questions underlying our study is whether students learning about a small marine organism like a symbion with multiple reproductive forms, or learning about a bird with frequent same-sex pairings can translate into a reduction of students’ negative attitudes and beliefs about humans who do not conform to normed sexual and gender roles. Our significant results indicate that many students readily made this leap between what is true about the biology of nonhuman life and what they saw as natural and acceptable for humans. This compares to a recent study on queer undergraduates students, in which Eddy et al. (2026) found that some students thought nonhuman examples of sex and gender diversity were impactful and transferable to influencing beliefs about humans, especially if the content was implemented as a consistent framing rather than one-off facts. However, in our interviews some students mentioned their belief in a separation between humans and animals. Previous research has indicated that the way youth think about “natural” categories of animals is related to the way they think about categories for humans like sex and gender (Rhodes & Gelman, 2009). The extent to which inclusive biology may impact a student’s views about humans may depend in part on how that student conceptualizes humans as a part of, an exception to, or entirely separate from, the rest of nature. Philosophers have long debated the validity and corollaries of the idea that humans are indeed animals, although the implications of this stance also depend on how one thinks about animals and their animal natures (Midgley, 1978). Some of the students in our study seemed to be grappling with these same questions, especially if faced with information about nonhuman species that contradicted their previous view of nature and/or human nature.

Relatedly, students may hold anti-LGBTQIA+ attitudes for reasons that have little to do with their understanding of biology or nature. Pseudoscientific beliefs are only one possible origin that can underly prejudicial attitudes, and exposure to new scientific information may do little to influence anti-LGBTQIA+ attitudes that are based in religious or cultural traditions (Hans et al., 2012). Furthermore, the authors acknowledge the potential pitfalls of inclusive biology education approaches that veer too far into the territory of the naturalistic fallacy—the idea that something being natural or present in nonhuman nature is what justifies its existence or makes it good (Fagundes & Coyne, 2023; Zemenick et al., 2022). However, we also argue that when “Biological Truth” and pseudoscientific justification are invoked for the directed suppression and erasure of LGBTQIA+ people, an appropriate response must necessarily include scientific counterevidence, and obligates the scientific community to become involved (Executive Order No. 14168, 2025; McLamore et al., 2023; Sedlacek et al. 2026). We also crafted our lesson plans with awareness of the very real legal constraints and repercussions that educators may face depending on the sociopolitical environment in which they are making teaching decisions, and for this reason it was important for at least some of the lesson plans to be free from any direct references to humans or human identities that would make them illegal in some U.S. States (Choi et al., 2025; Movement Advancement Project, 2022; Penney, 2022; Sadler et al., 2006).

### Inclusively and accurately defining biological sex

There are a wide array of perspectives on when or how to define biological sex, and these definitions will always be socially constructed as well as informed by science (Butler, 1999; Lane, 2009; Stryker, 2006). Many scientists attempt to be consistent with language that refers to sex cells as male if they are relatively smaller and refers to sex cells as female if the gametes are relatively larger (Heesch et al., 2021; G. A. Parker et al., 1972). This definition of gametes is favored by many scientists because it focuses on defining cells at a particular moment in time rather than classifying individuals, and can be applied across many different species, including plants that produce small male pollen and large female ovules, snails in which the same individual produces both male and female sex cells simultaneously, side-blotched lizards and harvest spiders in which there are multiple physically and behaviorally distinct morphs that produce the same type of sex cell, and in sequentially hermaphroditic species like many fish where an individual can change what type of sex cell it produces over its lifetime (Bachtrog et al., 2014; Painting et al., 2015; Pla et al., 2022; Sinervo & Lively, 1996). However, definitions of sexes can get murkier when a species has many mating types or reproductive forms like some fungi, protozoa, diatoms, and symbions (Burnett, 1965; Cervantes et al., 2013; Kaczmarska et al., 2013; Neves et al., 2010; Obst & Funch, 2003). In fact, due to the widely diverse arrangements of ingredients and mechanisms used in sexual reproduction across taxa, some argue there is no universal criteria for distinguishing females from males that can be universally applied for all eukaryotes (Gorelick et al., 2017). While it is beyond the scope of this article to attempt to provide a comprehensive consensus definition of biological sex, we do want to highlight our findings about the varied definitions held by the students in our study.

It is interesting to note that the LGBTQIA+ students’ we interviewed expressed understandings of the biology of sex that focused more on the function of gametes in a reproductive interaction, while the straight students we interviewed tended to describe definitions of sex using characteristics like genitalia, hormones, and behavior. In general, the content of our curriculum intervention attempted to present a view of biological sex aligned with recent biological research that does not assume sex is an immutable attribute of an individual, but rather a way to describe types of reproductive material and processes at one point in time. Although our interview sample was small, our findings suggest that LGBTQIA+ students’ understandings about biological sex may have already been more similar to a mutable and pluralistic view of biological sex than their straight peers before engaging with our curriculum intervention, which is also supported by the significant differences in quantitative scores we observed between LGBTQIA+ and straight students. We suggest that as scientists and educators, we continue to expand our own understandings of the nuanced spectrum of biological sex at the genetic, molecular, cellular, and phenotypic levels across many taxa, and that incorporating the latest scientific advancements into biology curriculum will also benefit the inclusion of LGBTQIA+ students by decreasing heteronormative, binary, and anthropocentric assumptions about the definitions and processes of biological sex (McLaughlin et al., 2023; Smiley et al., 2024).

Recent research suggests that gender and sex essentialism is present at a very young age in children (Pauker et al., 2020), although there are some intriguing differences in the way that cisgender and transgender children think about sex vs. gender and interpret terms like “boy” and “girl” (Gülgöz et al., 2021). We suggest that future research explore how LGBTQIA+ students form their understandings of the biology of sex and the role of their identities and lived experiences, and possibly their pursuit of different (outside school) sources of information than straight students. Regardless of the origin of these perspectives, our data demonstrate that students who are LGBTQIA+, or even those who have someone close to them who is LGBTQIA+, have differing attitudes and beliefs compared to straight students. This finding agrees with previous research on the Heteronormativity Attitudes and Beliefs Scale, which found differences in attitudes among straight and sexual minority participants (Habarth, 2015). Other research on essentialism more broadly has noted that an individual’s own identity can affect essentialist beliefs, but that there are complexities across different contexts such as race, sex, and ethnicity (Schudson & Gelman, 2023; Zhu & Scott, 2025). Similar to our findings, Schudson et al., (2019) found that transgender individuals had different and more complex definitions of gender and sex than cisgender individuals. It is vital to consider how these differing perspectives may differentially impact student experiences in STEM education, since outdated content may be particularly alienating to LGBTQ students in ways that it may not be for straight students. By changing the way we teach about the biology of sex, science education can be more inclusive while also being more scientifically accurate. Sticking with precise terminology that reflects current scientific knowledge of the wide variety of sexual systems that have evolved across many species, as others have also suggested (Cooper et al., 2020), avoids faulty generalizations that can veer into subjective cultural assumptions about human gender roles that can be especially alienating for girls and LGBTQIA+ science students.

#### LGBTQIA+ students are having distinct STEM education experiences that could contribute to disproportionate representation in STEM

There is mounting evidence of disparities in representation of LGBTQIA+ people in STEM majors and STEM careers (Freeman, 2020; Greathouse et al., 2018; Hughes, 2018). Survey research from STEM professional organization members shows that LGBTQIA+ professionals face systemic inequalities that lead to career limitations, harassment, and professional devaluation and higher intentions of leaving STEM (Cech & Waidzunas, 2021). Most students in our interviews saw no reason there should be differences in what subject a person was interested in based on their LGBTQIA+ identity. However, our qualitative findings explicitly point to the feeling of estrangement that LGBTQIA+ students can feel in “old-school” heteronormative science courses as a possible factor in quashing previous interest in STEM. The increased availability in recent years of other sources of information online likely means that some current LGBTQIA+ students are able to access updated and inclusive information on the biology of sex outside of school, and this access to information could increase their frustration and alienation when presented with outdated material in their science courses. Our efforts to understand STEM interest were hampered by issues with our survey instrument and we hope future research can find more reliable ways of exploring STEM interest to allow for comparisons among straight and LGBTQIA+ students, as well as any influence of inclusive curriculum interventions on STEM interest. It is possible that the STEM-CIS instrument was not reliable for high school students because of age-related differences, as it was originally developed with middle school students (Kier et al., 2014). We hope future research will continue to fill gaps in our knowledge about when and why LGBTQIA+ students leave STEM pathways, especially students with demonstrated prior interest in STEM.

We found differences among straight and LGBTQIA+ students in their initial attitudes and beliefs, as well as some differences in their responses to the curriculum intervention. Our findings suggest that straight students may have bigger changes in their beliefs in response to inclusive curriculum, however we note that broader impacts of inclusive curriculum for LGBTQIA+ students may be especially meaningful. As described in one of our interviews, the messages an LGBTQIA+ student receives in science education spaces can make them feel left out, or even like they themselves are not a relevant part of the natural world. Our findings are in alignment with Casper et al. (2022), who found that trans and gender nonconforming undergraduate students in biology courses experience feelings of erasure and exclusion.

Similarly, Beatty et al. (2021) found that students marginalized on the basis of race or ethnicity were especially likely to prefer ideologically aware science curriculum. School climate surveys also show that inclusive curriculum content significantly impacts how safe LGBTQIA+ students feel at school (Kosciw et al., 2013, 2022)

### Challenges and future directions

First, we acknowledge that there are limitations to a quasi-experimental longitudinal design and we cannot conclusively rule out other factors aside from our curriculum intervention that may have influenced the changes in attitudes and beliefs that we observed. For example, the research team was aware that some of our teacher partners were members of the LGBTQIA+ community themselves, and we do not have any data on whether students in our study were aware of or may have been impacted by the sociocultural identities of their instructors. However, we do believe several aspects of our design make it likely our curriculum intervention is the relevant cause of the observed shifts. For example, our sample included many participants who completed the activities independently and did not interact with any instructor, and further many of the participants took pre- and postsurveys on the same day, with little opportunity for factors other than the curriculum intervention to influence changes in their survey responses.

We also acknowledge that participant responses could have been impacted by social desirability bias, a phenomenon that could produce data that does not align with respondents’ true attitudes and beliefs. This study could be particularly vulnerable to this type of bias because social desirability bias is especially likely to exert influence in studies that address topics perceived to be socially or politically sensitive issues, and because some of the respondents in our study may have been aware of their teacher or interviewer’s social identities during their participation (Bispo Júnior, 2022; Grimm, 2010). While we cannot eliminate the potential influence of social desirability bias, we tried to mitigate these potential impacts by emphasizing during interviews that we were looking for opinions, and that there were no right or wrong answers. Social desirability bias was less likely to impact survey results since surveys were completed online while the participant was alone, and it was very clear the participants’ anonymity was ensured (Bispo Júnior, 2022).

There is also a possibility that there was self-selection bias in students who chose to take the survey (Berg, 2005). Some of the students within classrooms did not complete surveys because they did not have parental consent or because of absences from school, but it is also possible that some students who chose not to take the survey were more likely to hold certain attitudes, which could have resulted in a nonrepresentative sample of the student population.

However, research on underreporting and non-response bias has found that survey response rates tend to be higher for participants who hold both extreme positive and extreme negative views about the survey topic (Hu et al., 2017; Popovic & Huecker, 2026), which suggests this type of balanced response bias would not prevent us from drawing meaningful conclusions from our sample.

One recurring issue in our interviews was the lack of agreement between our research team and the participants on the use of the terms sex and gender. Despite our best efforts to be clear, and offer definitions in survey directions as recommended by the developers of the scales, our discussions in interviews showed that some students still considered sex and gender to be synonymous. This conflation is understandable given the widespread colloquial usage of the terms as interchangeable. Indeed, even for those who define sex and gender as distinct concepts, the impacts of sex and gender identities on people’s lived experiences can be difficult to separate because the two are so often intertwined (Viloria et al., 2020). Some have even suggested that attempting to treat sex and gender as fully distinct concepts for humans is futile (Fausto-Sterling, 2019; Lane, 2009; van Anders, 2015). We believe it is important to note this lack of clean separation of the two words and concepts in our data and consider the conflation of sex and gender to be both a challenge for our study and a finding in and of itself. The lack of shared vocabulary between researchers and interviewees can create difficulty in interpreting results from research instruments that use definitions that may be in tension with participant understandings, and we recommend that future researchers remain aware of this tension in design and analysis of future studies. For example, the transphobia scale in our study assumes a definition of gender as distinct from sex. If a participant reads the words gender and sex as synonymous, this prompt from item 9 of the Transphobia scale could be interpreted as a question about anatomical sex: “A person’s genitalia define what gender they are--a penis defines a person as being a boy, a vagina defines a person as being a girl.” It is possible that this linguistic issue could have contributed to the lack of significant impacts we observed on the transphobia scale.

Another challenge was our difficulty in recruiting teachers who were able to have their classrooms participate as research sites. While over 100 teachers contacted us to request inclusive teaching materials, most were unable to participate in data collection due to perceived or actual resistance from school administrators or their wider community. Our experiences conducting this study suggest that there is a great demand from educators for updated inclusive teaching materials, but as others have found, lack of curriculum resources is only one of several potential barriers that instructors may face (Driessen et al., 2024; Sadler et al., 2006). We also believe based on our interactions with educators while conducting this study that many teachers interested in inclusive science education may already be implementing interventions, but doing so quietly in order to avoid outside scrutiny of their teaching decisions. Relatedly, we allowed participating teachers the flexibility to choose from the four lessons what they were comfortable implementing to allow as many teachers as possible to participate, but this did prevent us from parsing out the impacts of specific lessons within the curriculum collection. We hope to home in on what specific content was most impactful in future studies, especially content that is explicitly about humans vs. content that explores the biology of sex through nonhuman examples.

We also believe much remains to be explored that arose during our student interviews that we were unable to pursue further. Our study timeline was also unexpectedly shortened, which prevented us from collecting more student interviews that would have provided greater richness to our data sample. In particular, we encourage future studies to target recruitment of more students with distinct LGBTQIA+ identities, especially transgender identities and also intersex identities. Even within research on sex and gender diversity in education, specifically transgender perspectives have sometimes been left behind (Airton & Koecher, 2019; Rende Mendoza & Johnson, 2024). The LGBTQIA+ umbrella encompasses many facets, and a larger sample would allow exploration of potential differing views between different identities within that umbrella. We also hope future research will explore connections between feminist and queer approaches to science education reform and the potential for interventions that simultaneously benefit girls and LGBTQIA+ students in STEM, as this seems like a natural pairing stemming from the multiple impacts of gender essentialism. Lastly, we suggest a future direction of research that focuses on the benefits of updated, GSD-inclusive science education on all science students, possibly through encouraging creative scientific hypothesis development, reducing deterministic and anthropocentric fallacies, and promoting authentic, connected learning.

### Implications

In the average science classroom today, there are likely several students who are LGBTQIA+, at least one student living with a caregiver who is not their genetic parent, many students who will never produce genetic offspring of their own for a variety of reasons, and depending on the size of the student population, one student at the school who is intersex (Blackless et al., 2000; Hemez et al., 2024; IPSOS, 2021; Minkin et al., 2024; MJ et al., 2018; Tierney & Cai, 2019).

Our call to educators is to think about how life science education can avoid assumptions that pretend these students are outside biological reality, or imply their existence is unnatural or unprecedented.

Our project has not only produced valuable research insights but also valuable curriculum tools towards achieving the goal of inclusive and scientifically accurate life science education.

Our findings have important implications for teacher education and science curriculum development as well as education research. We want to acknowledge that while it is the authors’ role as education researchers to critically examine curriculum and pedagogical approaches, we know there are many committed science educators who are simply teaching the way that they were taught with the limited and often outdated materials they have been provided. We also acknowledge that our inclusive science curriculum will always be a work in progress as new information and more precise interpretations of biology and inclusive education emerge. Our freely available curriculum offers classroom-tested, practical resources for educators who are interested in translating this research into application and want a place to start. The full curriculum with teacher guide, slide presentations, and student handouts is available for free, and future professional development opportunities for science educators will be listed on the SIUE STEM Center website https://stemcenter.siue.edu/.

## ACKNOWLEDGEMENTS

We thank all of the study participants who engaged with this program and contributed their thoughts to the study. We thank our teacher partners and curriculum reviewers Sam Long, Lewis Stellar, Thyme Masters, Laura Peña, Julian Laferrera, and Kirsten Milks. We are grateful to our advisory team Sharon Locke, Patrick Grzanka, Ren Rende Mendoza, and Harper Keenan, and to SIUE STEM Center office manager Dawn Olive. We thank Gwenn Seemel for providing artwork used in lesson materials. The study was partially funded by the National Science Foundation under award #2321243, which was prematurely terminated because the project no longer effectuated agency priorities. The remainder of the project was supported by the STEM Center and the Provost’s office of Southern Illinois University Edwardsville.

## Conflict of interest

The authors declare no financial conflict of interest.

## Notes

### Competing Interest Statement

The authors have declared no competing interest.

