## Supplemental files for "Inclusive Biology Curriculum Interventions Can Reduce High School Students’ Bioessentialist Beliefs"

Author preprint

#### Survey

*Note that participants could see the directions but not the title of each survey section. This shows the text from the post survey version which included the demographics questions.*

You will now answer the same survey questions you answered before the biology lessons. You should answer these survey questions based on what you think now-- it is okay if your answers are different or the same as your answers were before.

**STEM Interest**

Please mark how you feel about each statement. As you read the sentence, you will know whether you agree or disagree. Select the circle that describes how much you agree or disagree. The questions about your science class refer to the science class you are currently in.

1. I am able to get a good grade in my science class.
2. I am able to complete my science homework.
3. I plan to use science in my future career.
4. My parents would like it if I choose a science career.
5. I will work hard in my science classes.
6. If I do well in science classes, it will help my in my future career.
7. I am interested in careers that use science.
8. I like my science class.
9. I have a role model in a science career.
10. I would feel comfortable talking to people who work in science careers.
11. I know of someone in my family who uses science in their career.

Please use this space if you would like to explain why you did not answer or did not understand a particular question.

Author preprint

**Heteronormative Attitudes and Beliefs Scale**

(H) denoted heteronormativity subscale

(GE) denotes gender essentialism subscale

* denotes items that were reverse scored

Below are some statements representing different attitudes and beliefs. You will probably find that you agree with some of the statements, and disagree with others, to varying extents. Please indicate your reaction to each statement by clicking the appropriate circle next to each statement

1. In healthy intimate relationships, girls may sometimes take on stereotypical ‘male’ roles, and boys may sometimes take on stereotypical ‘female’ roles. *(H)
2. In intimate relationships, girls and boys take on roles according to gender for a reason; it’s really the best way to have a successful relationship. (H)
3. There are only two sexes: male and female. (GE)
4. People should partner with whomever they choose, regardless of sex or gender. *(H)
5. Gender is the same thing as sex. (GE)
6. Femininity and masculinity are determined by biological factors, such as genes and hormones, before birth. (GE)
7. All people are either male or female. (GE)
8. Things go better in intimate relationships if people act according to what is traditionally expected of their gender. (H)
9. Gender is a complicated issue, and it doesn’t always match up with biological sex. *(GE)
10. It’s perfectly okay for people to have intimate relationships with people of the same sex. *(H)
11. People who say that there are only two legitimate genders are mistaken. * (GE)
12. Gender is something we learn from society. * (GE)
13. There are particular ways that men should act and particular ways that women should act in relationships. (H)
14. The best way to raise a child is to have a mother and a father raise the child together. (H)
15. Sex is complex; in fact, there might even be more than 2 sexes. *(GE)
16. Women and men need not fall into stereotypical gender roles when in an intimate relationship. *(H)

Please use this space if you would like to explain why you did not answer or did not understand a particular question.

Author preprint

**Transphobia Scale**

Please mark how you feel about each statement. As you read the sentence, you will know whether you agree or disagree. Select the circle that describes how much you agree or disagree.

1. I don’t like it when someone is flirting with me, and I can't tell if they are a boy or a girl.
2. I think there is something wrong with a person who says that they are neither a boy nor a girl.
3. I would be upset, if someone I'd known a long time revealed to me that they used to be another gender.
4. I avoid people on the street whose gender is unclear to me.
5. When I meet someone, it is important for me to be able to identify them as a boy or a girl.
6. I believe that the male/female dichotomy is natural.
7. I am uncomfortable around people who don't conform to traditional gender roles—aggressive girls or emotional boys.
8. I believe that a person can never change their gender.
9. A person's genitalia define what gender they are-- a penis defines a person as being a boy, a vagina defines a person as being a girl.

Please use this space if you would like to explain why you did not answer or did not understand a particular question.

Author preprint

**Gender/sex Beliefs Scale**

Indicate your level of agreement with the following statements about gender and sex. Also, please note these definitions for terms some people might be unfamiliar with:

Transgender - a person whose gender identity is different from the gender they were assigned at birth. Example: "Michael is a transgender man. He was labeled a girl at birth and currently identifies as a man."

Cisgender - a person whose gender identity is the same as the gender they were assigned at birth. Example: "Alyssa is a cisgender woman. She was labeled a girl at birth and currently identifies as a woman."

Nonbinary - a person whose gender identity exists beyond woman or man or involves both. Non-binary identities include genderqueer, agender, etc. Example: "Taylor is non-binary. Taylor was labeled a boy at birth but is now agender, and does not identify with man or woman, or any gender."

1. A person's gender can change over time.
2. Nonbinary gender identities are valid.
3. Nonbinary gender identities have always existed.
4. People who express their gender in ways that go against society's norms are just being their true selves.
5. Gender is about how you express yourself (e.g., how you dress or act).
6. Being a girl or a boy has nothing to do with what genitals you have.
7. The only thing that determines whether someone truly is a girl or a boy is whether they identify as a girl or a boy.
8. Transgender identities are natural.
9. It would be best if society stopped labeling people based on whether they are female or male.
10. There are many different gender identities people can have.
11. Biological sex is not just female or male; there are many possibilities.
12. It is possible to have more than one gender identity at the same time.
13. Not all cultures have the same gender identities.

Please use this space if you would like to explain why you did not answer or did not understand a particular question.

Author preprint

**Demographic Questions**

Are you lesbian, gay, bisexual, transgender, queer, intersex, or asexual (LGBTQIA +)?
0 Yes
0 Maybe
0 No
0 I don’t understand the question.

Do you have an immediate family member, someone in your household, or a close friend who is lesbian, gay, bisexual, transgender, queer, intersex, or asexual (LGBTQIA +)?
0 Yes
0 Maybe
Q No
0 I don"t understand the question.

What is your current gender?
0 Boy/man
0 Girl/woman
0 Nonbinary
0 ___________Other
0 I don"t understand the question.

What sex was assigned on your original birth certificate?
O M
O F
O x
0 I don't understand the question.

Do you identify as transgender, nonbinary, gender expansive, Two-Spirit, and/ or gender non-conforming?
0 Yes
0 Maybe
0 No
0 I don't understand the question.

Are you intersex or do you have a difference of sexual development (DSD)?
0 Yes
0 Maybe
0 No
0 I don't understand the question.

What are your ethnic and/ or racial identities? Select ALL that apply and add any extra details in the box.
0 Mixed/Multiple Races

Author preprint

0 Black/ African American
0 White
0 Alaska Native/ American Indian
0 Asian
0 Middle Eastern/ Arab/North African
0 Latinx/Hispanic
0 Native Hawaiian/Other Pacific Islander
0 _____________ More details

If you live in the United States, in which state do you currently go to school?
What is your current grade level?
0 9
0 10
0 11
0 12
0 I am not in school because I graduated.

Author preprint

**Supplemental Table 1. Results of confirmatory factor analysis (CFA) for each of the five constructs we sought to measure in the student survey (N=127)**

| | $\chi^2$ | *df* | *p* | ratio $\chi^2$/*df* | CFI | RMSEA | SRMR |
|---|---|---|---|---|---|---|---|
| STEM interest | 133.4 | 44 | <.0001 | 3.03 | 0.734 | 0.136 | 0.117 |
| Heteronormative beliefs: normative behavior | 97.9 | 20 | <.0001 | 4.89 | 0.813 | 0.200 | 0.06 |
| Heteronormative beliefs: essential, binary sex and gender | 52.8 | 20 | <.0001 | 2.64 | 0.93 | 0.136 | 0.04 |
| Transphobia | 59.6 | 27 | <.0001 | 2.21 | 0.95 | 0.111 | 0.04 |
| Gender/Sex diversity affirmation | 134.9 | 65 | <.0001 | 2.07 | 0.93 | 0.122 | 0.03 |

We specified models for the CFA according to the structure proposed by the developers of each scale (Habarth 2015; Nagoshi et al. 2008; Schudson & van Anders, 2022). For the HABS, survey items were assigned to the appropriate subscale and some survey items were reverse scored as indicated by the developers of the scale (Habarth 2015). We used the robust maximum-likelihood estimation because of the ordinal nature of our data. Means values for the survey items ranged from 2.78 to 4.95, and survey items had no more than 3.1% missing values, apart from item 6 on the Transphobia scale as discussed in the manuscript. Nearly all items had skewness less than |1| and kurtosis less than |2|, and the only values outside these ranges were items from the STEM interest scale. We compared our CFA results against suggested cutoff values CFI ≥ 0.9, SRMR ≤ 0.08, and RMSEA ≤ 0.08, and prioritized the SRMR in our interpretation of the CFA output because it is more reliable at different sample sizes (Hu and Bentler 1999; Knekta et al. 2019; Marsh et al. 2004; Perry et al. 2015). We also report the ratio $\chi^2$/df because significance of the chi square test alone can be difficult to interpret, and others have suggested that a ratio less than 3 should increase confidence in the proposed model (Mousa et al. 2020). Based on our interpretation of the CFA we chose to exclude the STEM interest scale from our subsequent analyses and proceed with the other four scales.

Author preprint

**Supplemental Table 2. Factor loadings for each survey item from the CFA**

| Loadings STEM interest | Estimate | Std Error | Wald Z | Prob>\|Z\| |
|---|---|---|---|---|
| STEM interest → Stelar STEM-CIS_1 | 1 | 0 | . | . |
| STEM interest → Stelar STEM-CIS_2 | 0.2450466 | 0.2132602 | 1.1490501 | 0.2505 |
| STEM interest → Stelar STEM-CIS_3 | 4.0960555 | 1.8313575 | 2.2366226 | 0.0253* |
| STEM interest → Stelar STEM-CIS_4 | 2.058462 | 0.9501023 | 2.1665687 | 0.0303* |
| STEM interest → Stelar STEM-CIS_5 | 0.6334628 | 0.2555946 | 2.478389 | 0.0132* |
| STEM interest → Stelar STEM-CIS_6 | 2.7528006 | 1.1831176 | 2.3267346 | 0.0200* |
| STEM interest → Stelar STEM-CIS_7 | 4.2013486 | 1.8829579 | 2.2312494 | 0.0257* |
| STEM interest → Stelar STEM-CIS_8 | 1.7602324 | 0.6727519 | 2.6164661 | 0.0089* |
| STEM interest → Stelar STEM-CIS_9 | 2.6699403 | 1.1873744 | 2.2486087 | 0.0245* |
| STEM interest → Stelar STEM-CIS_10 | 0.9230627 | 0.4713412 | 1.9583746 | 0.0502 |
| STEM interest → Stelar STEM-CIS_11 | 1.1029093 | 0.5881074 | 1.8753535 | 0.0607 |
| **Loadings HABS normative behavior** | **Estimate** | **Std Error** | **Wald Z** | **Prob>\|Z\|** |
| HABS norm → REV HABS1 | 1 | 0 | . | . |
| HABS norm → HABS_2 | 1.7689976 | 0.3218605 | 5.4961617 | <.0001* |
| HABS norm → REV HABS4 | 1.8045432 | 0.3416475 | 5.2818869 | <.0001* |
| HABS norm → HABS_8 | 1.8412157 | 0.3191355 | 5.7693859 | <.0001* |
| HABS norm → REV HABS10 | 2.0342267 | 0.3480995 | 5.843809 | <.0001* |
| HABS norm → HABS_13 | 1.8974674 | 0.3603377 | 5.2658034 | <.0001* |
| HABS norm → HABS_14 | 1.689433 | 0.3116997 | 5.4200658 | <.0001* |
| HABS norm → REV HABS 16 | 1.0919175 | 0.2135712 | 5.1126636 | <.0001* |

Author preprint

| Loadings HABS essential, binary sex and gender | Estimate | Std Error | Wald Z | Prob>\|Z\| |
|---|---|---|---|---|
| HABS Essen → HABS_3 | 1 | 0 | . | . |
| HABS Essen → HABS_5 | 1.0456444 | 0.1095275 | 9.5468672 | <.0001* |
| HABS Essen → HABS_6 | 0.6729473 | 0.0870202 | 7.7332285 | <.0001* |
| HABS Essen → HABS_7 | 1.1262905 | 0.0678979 | 16.588013 | <.0001* |
| HABS Essen → REV HABS9 | 1.061926 | 0.1156032 | 9.1859531 | <.0001* |
| HABS Essen → REV HABS11 | 0.970799 | 0.1059537 | 9.1624793 | <.0001* |
| HABS Essen → REV HABS12 | 0.8656359 | 0.1093229 | 7.9181598 | <.0001* |
| HABS Essen → REV HABS 15 | 1.0502165 | 0.0818397 | 12.832607 | <.0001* |

| Loadings Transphobia | Estimate | Std Error | Wald Z | Prob>\|Z\| |
|---|---|---|---|---|
| Transphobia → Transphobia Scale_1 | 1 | 0 | . | . |
| Transphobia → Transphobia Scale_2 | 1.4263098 | 0.1620625 | 8.8009835 | <.0001* |
| Transphobia → Transphobia Scale_3 | 1.3097408 | 0.1524505 | 8.5912556 | <.0001* |
| Transphobia → Transphobia Scale_4 | 0.9694106 | 0.1385586 | 6.9963953 | <.0001* |
| Transphobia → Transphobia Scale_5 | 1.3113168 | 0.1489686 | 8.8026386 | <.0001* |
| Transphobia → Transphobia Scale_6 | 0.6976171 | 0.1467049 | 4.7552407 | <.0001* |
| Transphobia → Transphobia Scale_7 | 0.8334826 | 0.1325303 | 6.2889971 | <.0001* |
| Transphobia → Transphobia Scale_8 | 1.4353079 | 0.1755235 | 8.1772968 | <.0001* |
| Transphobia → Transphobia Scale_9 | 1.4052043 | 0.1697272 | 8.2791913 | <.0001* |

Author preprint

| Loadings Gender/Sex diversity affirmation | Estimate | Std Error | Wald Z | Prob>\|Z\| |
|---|---|---|---|---|
| Affirm Divers → Affirmation Subscal_1 | 1 | 0 | . | . |
| Affirm Divers → Affirmation Subscal_2 | 1.0733252 | 0.0561664 | 19.109731 | <.0001* |
| Affirm Divers → Affirmation Subscal_3 | 0.9889661 | 0.0661371 | 14.953266 | <.0001* |
| Affirm Divers → Affirmation Subscal_4 | 0.9475352 | 0.0605802 | 15.641004 | <.0001* |
| Affirm Divers → Affirmation Subscal_5 | 0.8383419 | 0.0665258 | 12.601755 | <.0001* |
| Affirm Divers → Affirmation Subscal_6 | 0.9986078 | 0.0632814 | 15.780434 | <.0001* |
| Affirm Divers → Affirmation Subscal_7 | 1.0407593 | 0.0515052 | 20.206897 | <.0001* |
| Affirm Divers → Affirmation Subscal_8 | 1.0452409 | 0.0551411 | 18.955756 | <.0001* |
| Affirm Divers → Affirmation Subscal_9 | 0.865992 | 0.0766826 | 11.293196 | <.0001* |
| Affirm Divers → Affirmation Subscal_10 | 1.1096091 | 0.0493059 | 22.504579 | <.0001* |
| Affirm Divers → Affirmation Subscal_11 | 0.9193324 | 0.0779321 | 11.796576 | <.0001* |
| Affirm Divers → Affirmation Subscal_12 | 1.0198549 | 0.0558436 | 18.262709 | <.0001* |
| Affirm Divers → Affirmation Subscal_13 | 0.8431706 | 0.0758004 | 11.123566 | <.0001* |

Author preprint

**Interview Guide**

Remember, during this interview there are no right or wrong answers. We just want to know what you think, whatever that is. Having your full, honest answers is what helps our study.

- Tell me what you remember about the activities you participated in. Was there anything that stood out to you that you never knew before?
- Was there anything in the lesson plans that you had already learned or were familiar with?
  - Did you learn about those topics in school or elsewhere?
- How would you define biological sex in your own words? (remember there is no right answer here, we just want your perspective)
- Did the activities you participated in change the way you thought about the biology of sex?
- Some people would say that in nature, there can only be two sexes, male and female. What do you think about that?
- As you participated in the biology activities and the survey, what did you not understand?
- When you participated in this study, you took the survey, participated in the biology activities, and then later answered the same survey questions again. Do you think any of your answers changed on the survey between the first and the second time you answered the questions?
  - If so, why do you think your answers changed?
- Do you think that these biology activities would be a welcome addition in the science classes at your school? Why or why not?
- Do you feel like the science classes at your school have been welcoming to LGBTQ students?
- Do you think LGBTQ students are more or less likely to go on to study science at university or have jobs in science? Why do you think that is?
- What else do you want to say?

Author preprint

### Codebook

| Code Category | Codes | Subcodes | Description |
|---|---|---|---|
| Epistemology of Sex and Gender | Biological sex | at birth/born with | Biological sex begins or is assigned at birth |
| | | chromosomes | Biological sex is defined by chromosomes |
| | | genitalia | Biological sex is defined by external genitalia |
| | | hormones | Biological sex is defined/shaped by hormones |
| | | innate behavior | Biological sex is characterized by innate behavioral differences |
| | | role in reproduction | Biological sex is defined by role in reproduction |
| | | cells produced | Biological sex is defined by what sex cells are produced |
| | Binarism | Binary | Sex/gender are two mutually exclusive categories, includes references to male/ female |
| | | Nonbinary | Pluralistic, nonbinary view of sex or gender |
| | Heteronormativity | | Centers heterosexual reproductive activity, or social norms |
| | Conflation of sex and gender | | Interpretation of question or phrasing of response that conflates sex and gender as synonymous, or replaces one with the other |
| | Humans and nonhumans | | Compares and contrasts humans and nonhumans in sex/gender concepts or characteristics, or in science education content |
| | Immutability of sex and gender | gender can change | Gender can change in an individual over time |
| | | sex can change | Sex can change in an individual over time |
| | | sex is fixed | Sex cannot change in an individual over time |
| Impacts of Curriculum Intervention | | | |
| | Attitudes and beliefs over time | changed following activities | Explicitly names that their thinking changed following the activities |

Author preprint

| Code Category | Codes | Subcodes | Description |
| --- | --- | --- | --- |
| | | did not change | States their thinking has not changed |
| | Feedback on curriculum | Positive | Positive feedback on the activities |
| | | Negative | Negative feedback on the activities |
| | Comparison to prior knowledge | had learned the info before | The student was familiar with the information in the activities |
| | | had not learned the info before | The information in the activities was new |
| Education in Social Context | Sociocultural identity in science | | Sociocultural identity matters in science, may include implicit or explicit examples |
| | LGBTQ+ students in STEM | | The interviewees perspective on LGBTQ+ students in STEM spaces generally or in the classroom specifically |
| | Openness to inclusive education | | Support for the implementation of inclusive education practices |
| | Science Interest | | Interest, excitement, wonder, motivation about science |
| | School climate | | The interviewees perspective on whether educational environment is welcoming and inclusive for LGBTQ+ students |
| | Sociopolitical context | | Explicit acknowledgment or awareness of the larger sociopolitical context in which education takes place |

Author preprint
